# Climate change and reseeding shape richness-evenness relationships in a subalpine grassland experiment

**DOI:** 10.1101/2024.11.11.622915

**Authors:** Lina K. Mühlbauer, Andreas Klingler, Lukas Gaier, Andreas Schaumberger, Adam Thomas Clark

## Abstract

Grasslands face an uncertain future due to climate change. Although there is increased interest in the interdependencies of different biodiversity components, the effects of climate change on these relationships remain understudied. One of these is the richness-evenness relationship (RER), which is sensitive to altered species abundances in relation to richness. This relationship may be important as evenness and richness jointly shape diverse ecosystem functions, such as stability and productivity. As evenness affects productivity differently in low and high richness communities, the richness-evenness relationship is important to investigate, especially under climate change.

Here, we assess the effects of increased CO_2_ concentrations, temperature, and drought on the RER in a subalpine long-term (2010 - ongoing) grassland climate change experiment, and test whether these effects can be buffered by reseeding. We provide evidence that climate change alters the RER in our experiment, and that these changes occur independently of changes in richness and evenness separately. Reseeding erases the differences in RER between treatments and controls but fails to restore the negative RER initially found in controls. Further, we show that the dominant grass species in our system *(Arrhenatherum elatius)* responds differently to each climate change factor, with opposite effects in high vs. low richness plots, thereby largely determining the direction of the RER.

These results suggest that the RER can reveal additional insights on community responses to climate change and represents a different signal than evenness or richness alone. A more nuanced approach integrating evenness and maximizing richness in seed mixtures could be an important step forward to better match restoration treatments to particular community types and global change drivers.

## Introduction

Grasslands face an uncertain future due to climate change (Stevens et al. 2022). Climate change experiments suggest that biodiversity, productivity, and stability can be altered by increased temperature levels (Kardol et al. 2010; Wang et al. 2020; Fussmann et al. 2014), but elevation of CO_2_ concentrations could potentially ameliorate these effects (Reyes-Fox et al. 2014; Roy et al. 2016). Overall, warming seems to decrease richness and slightly also evenness, probably due to heat- tolerant species increasing in cover and susceptible species disappearing. An increase in atmospheric CO_2_ alters resource and water availability in the soil and therefore changes the competitive interactions between plant species. Consequently, the response to CO_2_ is suggested to be largely mediated by species identity (Niklaus et al., 2001, Potvin et al. 2007; Navas et al. 2002; Ramsier et al. 2005). In natural communities, CO_2_ seems to have no effect on species richness, but enhances species evenness in an experiment in the Swiss Alps (Niklaus et al., 2001). In a field experiment in Quebec in a less species-rich community, species loss and increases in species evenness were reported (Potvin and Vasseur 1997).

Also, elevated CO_2_ seems to reduce species loss driven by nitrogen addition, at least in low richness communities (Reich, 2009).

While many studies have quantified the effects of increased temperature and CO_2_ separately, evidence from controlled experimental settings with interactive effects is rarer, but they seem to range from simple and additive to highly complex interactions (Stevens et al. 2022). In most studies in temperate regions, species richness remains unchanged or even increases, while evenness decreases (Kardol et al. 2010; Liu et al. 2018) when grasslands are exposed to enhanced temperatures and CO_2_ concentrations. On the other hand, some studies suggest a dramatic loss of species richness (Zavaleta et al. 2003). Drought impact is expected to result in species loss, either through heat stress (Klein et al. 2004) or decreased soil moisture (White et al. 2014). In general, in grassland climate change experiments, biodiversity responses differ widely, especially depending on region (Bütof et al. 2012). This is especially true for drought responses, where the local climate and ecosystem properties, like community composition, determine the magnitude of drought effects (Isbell et al. 2015; Boeck et al. 2016; Gruner et al. 2017), and nutrient addition can amplify the impacts of drought (van Sundert et al. 2021). Additionally, biodiversity responses seem to lag behind to effects of climate change, especially in mountainous regions (Alexander et al. 2018; Rumpf et al. 2019).

Although there is increased interest in the interdependencies of different biodiversity components (e.g. Blowes et al. 2022) and the effects of climate change on the relationships between different biodiversity components remain understudied. One example of these interdependencies is the relationship between species richness and evenness over several communities, known as the “richness-evenness relationship” (RER). The RER could be an early sign of community changes, evident long before serious impacts such as extinction events take place, as proposed for evenness (Chapin et al. 2000). As the RER has been shown to vary in space, time, and across environmental conditions or trophic levels (Soininen et al. 2012; Zhang et al. 2012), a change in RER is likely an important indicator of underlying biological processes. Also, evenness and richness jointly shape diverse ecosystem functions, like productivity (e.g., Clare et al. 2022), as well as the stability of ecosystems (Wang et al. 2021).

Recently, Hordijk et al. (2023) were able to show that evenness mediates the global species richness- productivity relationship, such that at high species richness, even communities are more productive than uneven ones, whereas the contrary is true for low richness systems. This finding emphasizes the importance of investigating the relationship of evenness and richness, as changes in the relationship could indicate that less productive communities (low richness, high evenness and high richness, low evenness) get replaced by more productive communities (low richness, low evenness and high richness, high evenness) (Fig. 1, b). Stirling and Wilsey (2001) conducted regression analyses of published data on the relation between evenness and species richness and concluded that the RER is mostly positive in animals and negative in plants and fungi, while more comprehensive analyses are still missing. Of major importance in analyzing the RER is that many of the commonly used evenness metrics (such as Shannon’s or Pieloús evenness) are mathematically constrained by species richness, such that they necessarily increase or decrease with richness, independent of any variation in species composition (Smith und Wilson 1996; Tuomisto, 2012; Hordijk et al. 2023). As we examine richness- evenness relationships, the most important feature for an evenness metric in our analysis is species richness-independence – in other words, an effective evenness metric should be able to span its full range of possible values at any level of species richness. Studies investigating RERs use species richness independent evenness metrics (e.g., Hordijk et al., 2023) and evenness metrics that are constrained by species richness (e.g., Soininen et al., 2012). Here, we use the unbiased Gini coefficient (Lorenz 1905) commonly used to assess inequality in economics, which has been shown in previous studies to be a robust, species richness-independent evenness metric (Chao und Ricotta 2019).

**Figure 1:**
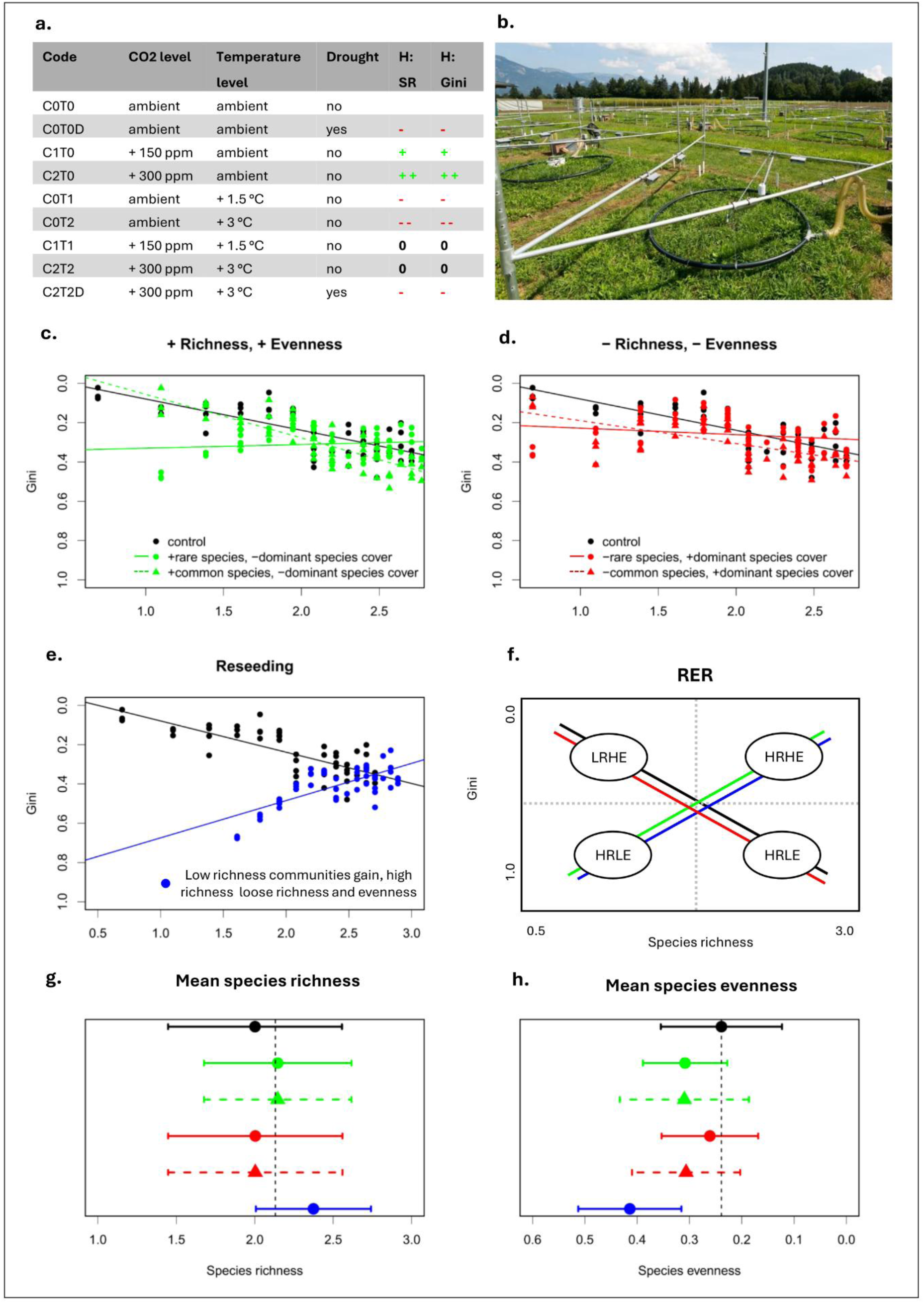
Conceptual figure showing hypotheses of potential kinds of biodiversity change, and their corresponding potential impacts on the RER in response to climate change (increase in CO_2_, temperature, and drought via rainout shelter) and reseeding based on simulated data (see Methods). Hypotheses for the effects on evenness and richness separately for each treatment (a., g., and h.) and their combined effect on RER (c., d., and e.) are shown. Colors indicate expected changes in biodiversity due to experimental treatments. Green indicates increases and red decreases in richness and evenness simultaneously. Black indicates control plots. We expect an increased species richness and evenness with higher CO_2_ concentrations, but an unchanged relationship between evenness and richness. The temperature treatment could lead to decreased richness and evenness, due to heat stress and reduced water availability. Simultaneous increase of CO_2_ and temperature could lead to unchanged evenness and richness and a negative RER, as CO_2_ could ameliorate the effects of temperature. With additional drought application, we expect species richness and evenness to decrease, as water and heat stress increase and are too high to be buffered by CO_2_ and therefore reduced evapotranspiration (**a.**). We expect no change in the direction of RER, as we expect the treatments to affect low and high richness communities in the same way. In controls, we expect higher diversity plots also tend to be dominated by a single abundant species (red triangles), with less abundance in low diversity plots leading to low evenness, as observed for natural plant communities and consequently to a negative RER (**c., d., e., and f.**). We expect the RER to change in direction, due to increases in richness and evenness, if the added species is rare, i.e., low percentage cover, and due to reseeding (**c.** and **e.**). We expect no change in the direction of the RER following decreases in richness and evenness, only changes in the steepness of the slope (**d).** When low-richness control plots are reseeded, several new species can establish, leading to an increase in richness and consistent evenness. In contrast, when high-richness plots are reseeded with a mixture containing a large amount of the already dominant species, the dominant species can competitively suppress other species in the mixture, thereby leading to decreased richness and slightly decreased evenness and therefore a change in the direction of the RER (**e.**). These changes, i.e. the replacement of low richness, high evenness (LRHE) and high richness, low evenness communities (HRLE) by low richness, low evenness (LRLE) and high richness, high evenness (HRHE) communities has consequences for productivity and stability (**f.**) and are only partly detectable by changes in mean (points and triangles) and standard deviation (lines) of richness and evenness separately (**g.** and **h.**). In controls, higher diversity plots also tend to be dominated by a single abundant species (red triangles), with less abundance in low diversity plots leading to low evenness, as observed for natural plant communities. b. shows one plot of the ClimGrass experiment in Raumberg-Gumpenstein, Styria, Austria (47°29’38.5" N 14°06’02.6" E, 732 m) after mowing, with the black fumigation ring and the heaters visible. Methods (1368)

In addition to the assessment of climate change effects on biodiversity, the mitigation of these effects increases in importance. Temperate grasslands in Europe contain the most diverse plant communities in Europe (WallisDeVries et al. 2002). Besides their biodiversity, European grasslands are economically important as a source of fodder, and, to a lesser extent, fiber, and energy. Consequently, central European grasslands are often intensively fertilized and mowed up to 6 times a year.

Intensively managed permanent agricultural grasslands are less diverse than extensively managed grasslands, with lower mowing frequencies and fertilizer application, but account for 34% of agriculturally used land in Europe (Eurostat 2020). Understanding the effects of climate change and investigating potential restoration strategies is therefore particularly relevant in these agricultural grasslands, due to the wide range of services and economic benefits they provide.

In grasslands, the most common restoration method is reseeding, independent of the kind of disturbance. There is also evidence that passive restoration, i.e. the recovery of natural grasslands without additional actions like reseeding, fails to completely restore biodiversity (Ladouceur et al. 2023). Evidence for the results of reseeding as a restoration strategy is mixed but has thus far been assessed primarily for soil properties, phenology, and productivity (Zhu et al., 2020; Wang et al., 2020).

How reseeding affects different facets of biodiversity, and their interdependencies, is often not assessed, as most conservation studies concentrate on boosting rare species abundances (Samson and Knopf 1994) or focus on creating or restoring species-rich, extensively used grasslands (Slodowicz et al. 2023). Therefore, an open and important question is whether and how 1) climate change affects the relationship between evenness and richness (RER) and 2) if reseeding can erase, or at least partially alleviate, these effects in intensively used grasslands.

Here, we test the effects of increased CO_2_ concentrations, temperature, and drought interactively on richness and evenness and on the RER in a unique subalpine, agricultural long-term (2010 - ongoing) climate change experiment and assess the effects of reseeding under climate change. We expect an increase in species richness and evenness with higher CO_2_ levels, as CO_2_ could ameliorate the effects of fertilization in the nutrient-rich grasslands in our experiment and enable less competitive species to colonize. The increase in temperature could lead to decreased richness and evenness, due to heat stress or decreased water availability. Simultaneous increase of CO_2_ and temperature would lead to unchanged evenness and richness, as CO_2_ could ameliorate the effects of temperature due to reduced evapotranspiration, thereby buffering the effects of increased temperature (Veronika Slawitsch et al. 2019). With additional drought application, we expect species richness and evenness to decrease, as water and heat stress increase and are too high to be buffered by CO_2_ and therefore reduced evapotranspiration (Fig. 1, a). Changes in species evenness alone lead to no changes in the RER, while the effect of species richness changes (addition or removal of species) depends on the abundance of the respective species. If a treatment leads to increased evenness and richness, we expect no change in RER if the added species is common, but if a rare species is added, we expect the RER to become positive (Fig. 1, c.). On the contrary, if a treatment is expected to decrease evenness and richness, we expect changes in the steepness of the slope, but no change in the direction of the RER. We expect a change in direction of RER by reseeding, if low richness, high evenness plots are reseeded, several new species can establish, leading to an increase in richness and consistent evenness. In contrast, when high-richness, low evenness plots are reseeded with a mixture containing a large amount of the already dominant species, the dominant species can competitively suppress other species in the mixture, thereby leading to decreased richness and slightly decreased evenness.

This leads to positive RER (Fig. 1, b. and e.). Additionally, we expect the RER to change independently from changes in richness and evenness separately (Fig. 1, f). These changes, i.e. the replacement of low richness, high evenness (LRHE) and high richness, low evenness communities (HRLE) by low richness, low evenness (LRLE) and high richness, high evenness (HRHE) communities have consequences for productivity and stability and underline the importance of investigating the effects of climate change not only on species richness and evenness, but also on their relationship.

### Experimental design

The *ClimGrass* climate change experiment was established at the Agricultural Research and Education Centre (AREC) in Raumberg-Gumpenstein, Styria, Austria (47°29’38.5" N 14°06’02.6" E, 732 m). This experiment focuses on agriculturally relevant management methods (e.g., fertilization, mowing, reseeding), and because of its subalpine location (Pötsch et al. 2020). With a mean annual precipitation of 1077 mm per year and a mean annual temperature of 8.5 °C, the location is representative of most low-elevation parts of the eastern European Alps. Climate diagrams for the changes in precipitation and temperature over the last decades can be found in the supplement (Fig. S1). The soil is mainly brown soil with a loamy texture consisting of 44.2% sand, 47.6% silt, and 8.3% clay with a mean pH of 5.7 (Herndl et al. 2011). The grassland was established in 2007 and seeded with the *DWB* Seed mixture (see S1, Tab. 4 for details) on bare soil; the same mixture was used for reseeding. The climate change treatments were started in 2010 and are still ongoing. The site can be classified as a nutrient-rich meadow that primarily consists of three C_3_ grass species: tall oat-grass (*Arrhenatherum elatius* L.), Kentucky bluegrass (*Poa pratensis* L.), and orchard grass (*Dactylis glomerata* L.), together with common forbs and legumes, e.g., dandelion (*Taraxacum officinale* L.) or red clover (*Trifolium pratense* L.).

The meadow is mowed three times per year during the growing season (April to October). Mineral fertilizer is also applied three times annually, resulting in a total amount of 90 kg N ha−1 y−1, 65 kg P ha−1 y−1, and 170 kg K ha−1 y-1 (Deltedesco et al. 2019). The field experiment is designed following a response surface approach (Piepho et al. 2017). It consists of 54 plots with a size of 16 m^2^ each. Experimental treatments impose various levels of CO_2_ (ambient, +150, +300 ppm; denoted by C0, C1, and C2), air temperature (ambient, +1.5°, +3 °C; denoted by T0, T1 and T2), and simulated drought (denoted by “D”). Interactive treatments between temperature, CO_2_ and drought are also implemented. The C2T2 treatment (+300 ppm and +3°C) is based on the most likely future climate change scenario for the European alps (Gobiet et al. 2014). All treatment descriptions can be found in Tab.1.

The temperature treatments are heated by six infrared radiators. The CO_2_ treatments are maintained by mini-FACE (Free Air Carbon Dioxide Enrichment) rings around the plots, where the CO_2_ concentration is controlled via a sensor in the plot center (Herndl et al. 2011). The temperature increase is applied all year round when the snow cover is below 10 cm, while CO_2_ is only added during daytime in the growing season. Control plots are also equipped with mini-FACE rings with ambient air flow and unconnected radiators to control for shading and unintended equipment effects (Herndl et al. 2011; Pötsch et al. 2020). Drought treatments with rainout shelters have also been included since 2017 (Veronika Slawitsch et al. 2019). These were used to simulate the effects of severe droughts, with a total reduction in incoming precipitation in 2017, 2019, 2020, 2021 and 2022. To end each drought period, drought treatments received 40 mm of irrigation supplied from collected rainwater, manually applied to the drought plots (see S1, Tab. S7). As an agricultural experiment, plots were periodically reseeded to help increase cover, after visual assessment for ants, voles, or cockchafer grubs. Reseeding took place in 2018, 2020 and 2021, mimicking the management methods that farmers in the area would have implemented. Because the reseeding mixture (see S1, Tab. S4) is identical with the initial seed mixture, no new species were introduced, and relative seed amounts matched those for the initial conditions of the experiment. The colonization of different species could occur additionally. The number of reseeded plots per treatment can be found in Tab. 1. Vegetation surveys were conducted in 2013, 2017, 2019 and 2022 to record species-level absolute cover in each plot. Absolute cover and species richness estimates were performed in mid to end of May in the 1 m^2^ harvest ring in the plot center (Peratoner und Pötsch 2019).

**Table 1:** Analyzed treatments. , corresponding codes, number of plots, and number of analyzed plots (reseeded from 2018) for each treatment.

| Code | CO<sub>2</sub> level | Temperature level | Drought | #Plots | #Analysed<br>(reseeded) |
| --- | --- | --- | --- | --- | --- |
| **C0T0** | ambient | ambient | no | 16 | 13 |
| **C0T0D** | ambient | ambient | yes | 4 | 4 |
| **C1T0** | + 150 ppm | ambient | no | 4 | 3 |
| **C2T0** | + 300 ppm | ambient | no | 6 | 2 |
| **C0T1** | ambient | + 1.5 °C | no | 4 | 3 |
| **C0T2** | ambient | + 3 °C | no | 6 | 4 |
| **C1T1** | + 150 ppm | + 1.5 °C | no | 3 | 2 |
| **C2T2** | + 300 ppm | + 3 °C | no | 7 | 4 |
| **C2T2D** | + 300 ppm | + 3 °C | yes | 4 | 4 |

## Data simulation

To better explore our hypotheses of the effects of changes in evenness and richness separately on the relationship between them, we simulated species richness and corresponding abundance for each species. We simulated species richness from a uniform distribution between 2 and 15 species for 54 plots. We sampled the percentage cover abundance for each species from a normal distribution with a mean of 20% cover. To account for the negative RER found in the available literature (Soininen et al., 2012; Zhang et al., 2012; Hordijk et al., 2023), we sampled the abundance of plots with less than 8 species from a normal distribution with a lower standard deviation of 5 %, and communities with more than 8 species with a higher standard deviation of 15%. The resulting overall percentage cover was then normalized to 100% cover, and Gini and species richness were calculated (see Methods: Biodiversity metrics), as performed for the actual experimental survey data. To add a rare species to plots, we sampled from a uniform distribution with 1 to 5 % cover. To remove a rare species, we removed the rarest species of the community in plots with species richness greater than 2. To decrease or increase evenness, we added or removed 5% cover to or from the dominant species. To simulate our expectations for the biodiversity changes due to reseeding, we sampled three rare species (uniform distribution; 1 to 5 % cover) to add to plots with less than 8 species, and added 40% cover to the dominant species.

### Data analyses

All analyses were conducted using R 4.2.2. (R Core Team, 2022).

To prevent post-treatment bias in subsequent analyses, we only used the vegetation data from plots that were reseeded in 2018, 2020, and 2021. Thus, we exclude data from plots that were never reseeded (in total, 15 plots). Also, within individual analyses, we used either solely data from plots that had not yet been reseeded or solely data from plots that had already been reseeded (i.e., we never mixed data from pre- and post-reseeding within a single analysis). All analyzed treatments and the corresponding number of replicates can be found in Tab. 1.

### Biodiversity metrics

To examine the effects of experimental treatments on RER, species richness and evenness were calculated for each plot in each year. The unbiased Gini metric, calculated as:

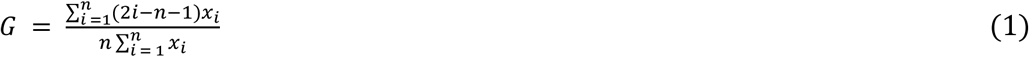

was used to quantify the richness independent evenness (Chao and Ricotta 2019), where x is the observed abundance in percentage cover of a species, n is the number of species observed, and i is the rank of values in ascending order (Buchan, 2002). We did so using the R package DescTools, version 0.99.46. Before calculating evenness, cover estimates were normalized to absolute (i.e., summing to 100 percent) cover (not counting bare ground). Importantly, note that Gini, in contrast to commonly used evenness metrics (e.g., Shannon’s or Pielou’s evenness), ranges between 0 and 1, where 0 indicates complete evenness (i.e., all species having equal abundance), and 1 indicates complete dominance by a single species. For easier interpretability, all evenness axes in the main figures are flipped, i.e., to display 1 at the bottom of the axis and 0 at the top. To investigate differences in species richness and evenness, we fit linear regressions predicting evenness and log-transformed species richness as a function of treatment. We included plot as a random effect, and year as a categorical fixed effect (i.e., resulting in models of the form gini/species richness ∼ Treatment + date_factor + (1|plot)). Differences in species richness and evenness between years can be found in the supplement (Fig. S6-10). All linear regressions were performed using Bayesian Generalized Linear Multivariate Multilevel Models in the brms R package, version 2.17.0. We used the default settings for the package, i.e., with 1000 burn-in steps, 1000 MCMC optimization steps, four chains, and flat priors (Bürkner 2017). In all cases, models achieved convergence, with R-hat estimates of <1.1 for all models and parameters. Significant differences in species richness and evenness between treatments and control were identified by comparing the posterior distributions of intercept estimates across the fitted regressions.

### Richness-evenness- relationships (RER)

To investigate differences in RER between control and treatments, we fit linear regressions predicting evenness as a function of logarithmic species richness, with separate regression fits for each combination of experimental treatments. Recall that we analyzed only plots that were reseeded at some point in 2018, 2020, and 2021, and that in all cases, we separately fit functions for the RER prior to reseeding, and after reseeding. All linear regressions were performed using Bayesian Generalized Linear Multivariate Multilevel Models in the brms R package, version 2.17.0. We used the default settings for the package, i.e., with 1000 burn-in steps, 1000 MCMC optimization steps, four chains, and flat priors. In all cases, models achieved excellent convergence, with R-hat estimates of <1.1 for all models and parameters. Survey year and treatments as factors were included as fixed effects in the models, whereas plot identity was modeled as a random effect acting on the model intercept, i.e., of the form (1|Plot) (Bürkner, 2017). Significant differences in the RER across experimental treatments were identified by comparing the posterior distributions of intercept and slope estimates across the fitted regressions.

#### A. elatius cover

To examine how the abundance of the dominant species *A. elatius* was associated with the difference in RER before and after reseeding, we calculated the difference in *A. elatius* cover before reseeding (vegetation surveys in 2013, 2017) vs. after reseeding (vegetation surveys in 2019, 2022), as well as differences in species richness and evenness. We then fitted regression lines for the relationship between log-transformed species richness vs. evenness, with corresponding confidence intervals, using brms. Log transformation was necessary to meet normality assumptions for model residuals.

## Results

### Treatments increase richness and reduce evenness

In general, experimental CO_2_, temperature, and drought treatments tended to increase richness and reduce evenness before reseeding, with two minor exceptions (Fig. 2, a-b). First, in the CO_2_ treatments at ambient temperature levels (C1T0, C2T0), mean evenness was somewhat higher than in control plots. Second, in the +1.5 °C treatments at ambient CO_2_ (C0T1), mean richness was somewhat lower than in control plots. Before reseeding, species richness was only increased in the Scenario 2 (+300 ppm, 3°C) treatment, while evenness was significantly decreased in both scenarios (C1T1: +150 ppm, 1.5°C; C2T2: +300 ppm, 3°C) (Tab. 2). The RER in the control treatment before reseeding is negative in 2013 and in 2017, whereas the differences between control and treatment RER is more pronounced in 2017 (Fig. 2, b. and c.) Therefor, we controlled for temporal variation by including year as a continuous fixed-effect predictor in regression analyses described below (Fig. 3 and 4) – recall again that all comparisons were conducted either between treatments before reseeding, or after reseeding, but never with mixed data from both time-periods.

**Figure 2:**
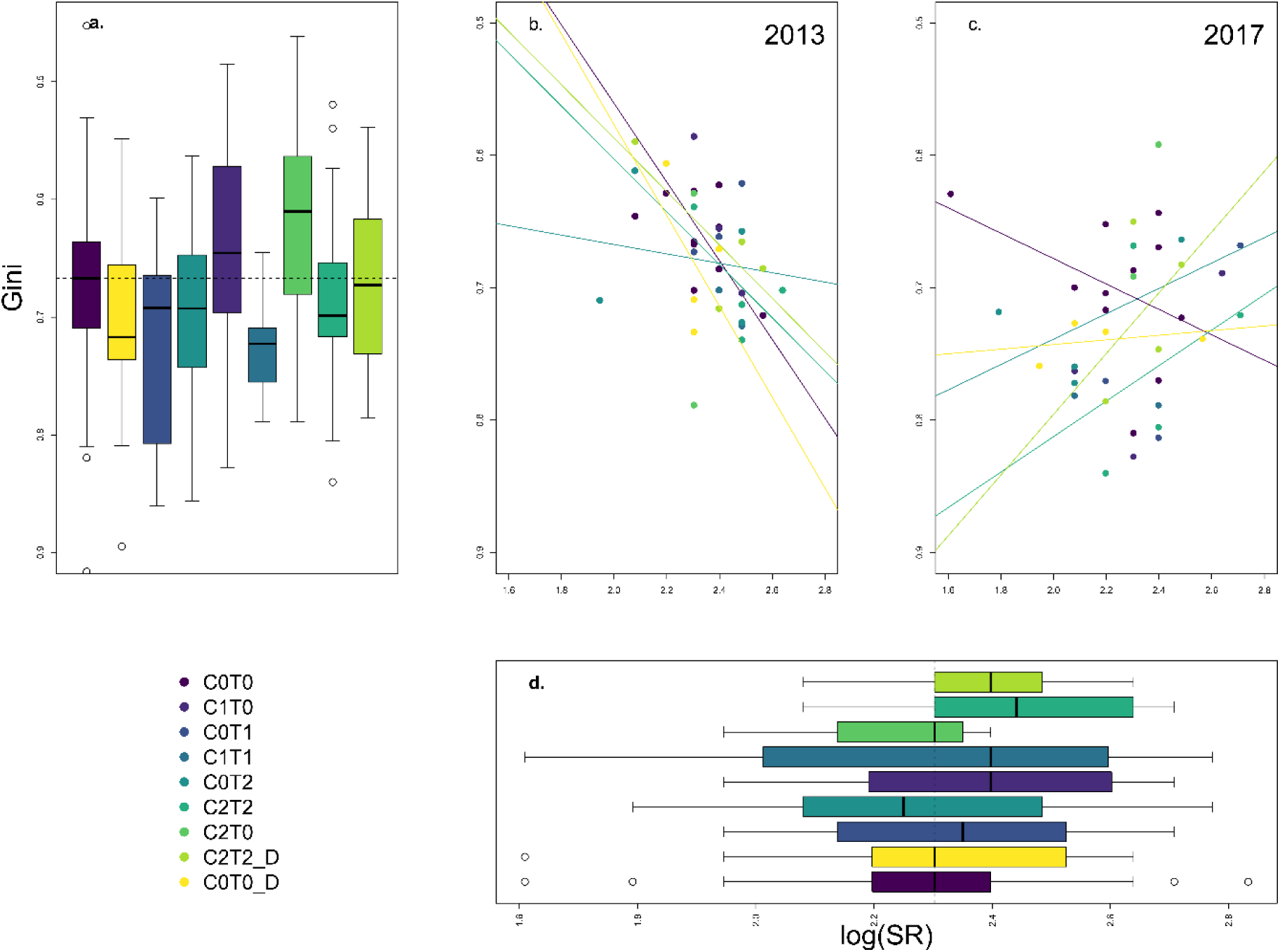
Treatment differences in **a.** mean evenness, c., and d. richness evenness relationship before reseeding (2013 and 2017) and **d.** mean species richness. Only results from prior to reseeding are shown, but only from plots that were eventually reseeded in 2018, i.e., from years 2013 and 2017. Dashed line in boxplots indicates value of the control (C0T0). Black bars indicate median, and boxes indicate 25% and 75% quantiles. Colors indicate treatments. Treatment codes indicate temperature (T1 = +1.5 °C, T2 = +3°C) and CO_2_ (C1 = +150 ppm, T2 = +300 ppm) levels and drought treatment (D = drought). Gini is used as an evenness metric, where 0 indicates complete evenness, and 1 indicates complete dominance. For easier interpretability, all evenness axes in the main figures are flipped, i.e., to display 1 at the bottom of the axis and 0 at the top. Data are from vegetation surveys conducted in 2013 and 2017, prior to reseeding. Note, however, that to avoid post-treatment biases, we only include data from plots that were eventually reseeded as part of the experiment.

**Figure 3:**
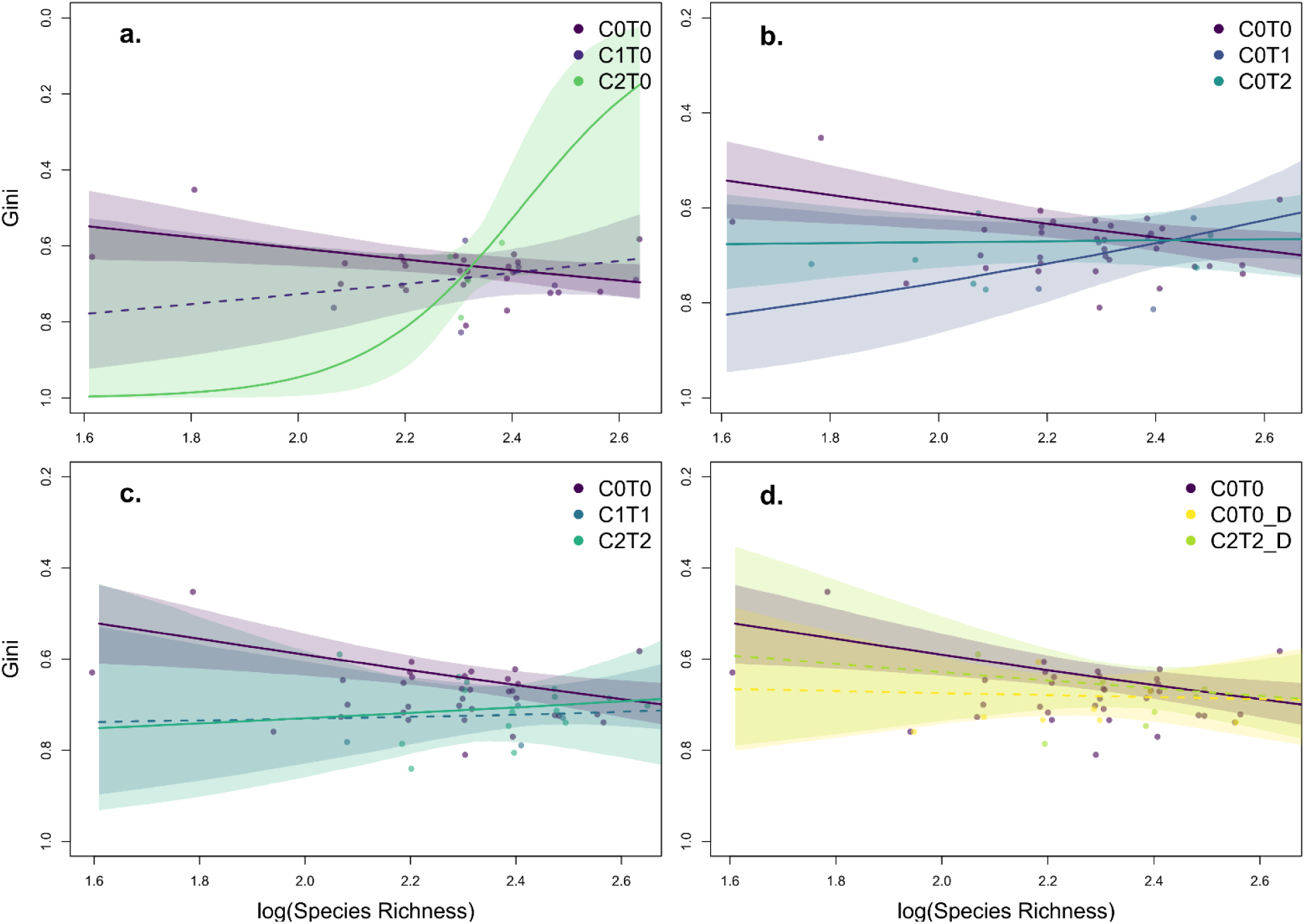
Climate change treatments alter richness-evenness relationships (RER) prior to re-seeding for control plots vs. **a.** CO_2_ treatments (C1T0: +150 ppm and C2T0: +300 ppm), **b.** temperature treatments (C0T1: +1.5 °C and C0T2: +3°C), **c.** climate change scenario 1 (C1T1: +150 ppm, + 1.5°C) and 2 (C2T2: +300 ppm and +3°C), and **d.** drought treatments C0T0_D, C2T2_D (implemented only for control and scenario 2 plots). Shaded regions show 95% confidence intervals. Points show single plots; points and lines are colored by treatment level (purple for control, blue for first level, and yellow for second level). Dashed lines indicate treatments with slopes not significantly different from the control (see Fig. 2). Regression structure is described in the data analyses section of the methods. Gini is used as an evenness metric, where 0 indicates complete evenness, and 1 indicates complete dominance. For easier interpretability, all evenness axes in the main figures are flipped, i.e., to display 1 at the bottom of the axis and 0 at the top. Data are from vegetation surveys conducted in 2013 and 2017, prior to reseeding. Note, however, that to avoid post- treatment biases, we only include data from plots that were eventually reseeded as part of the experiment.

**Figure 4:**
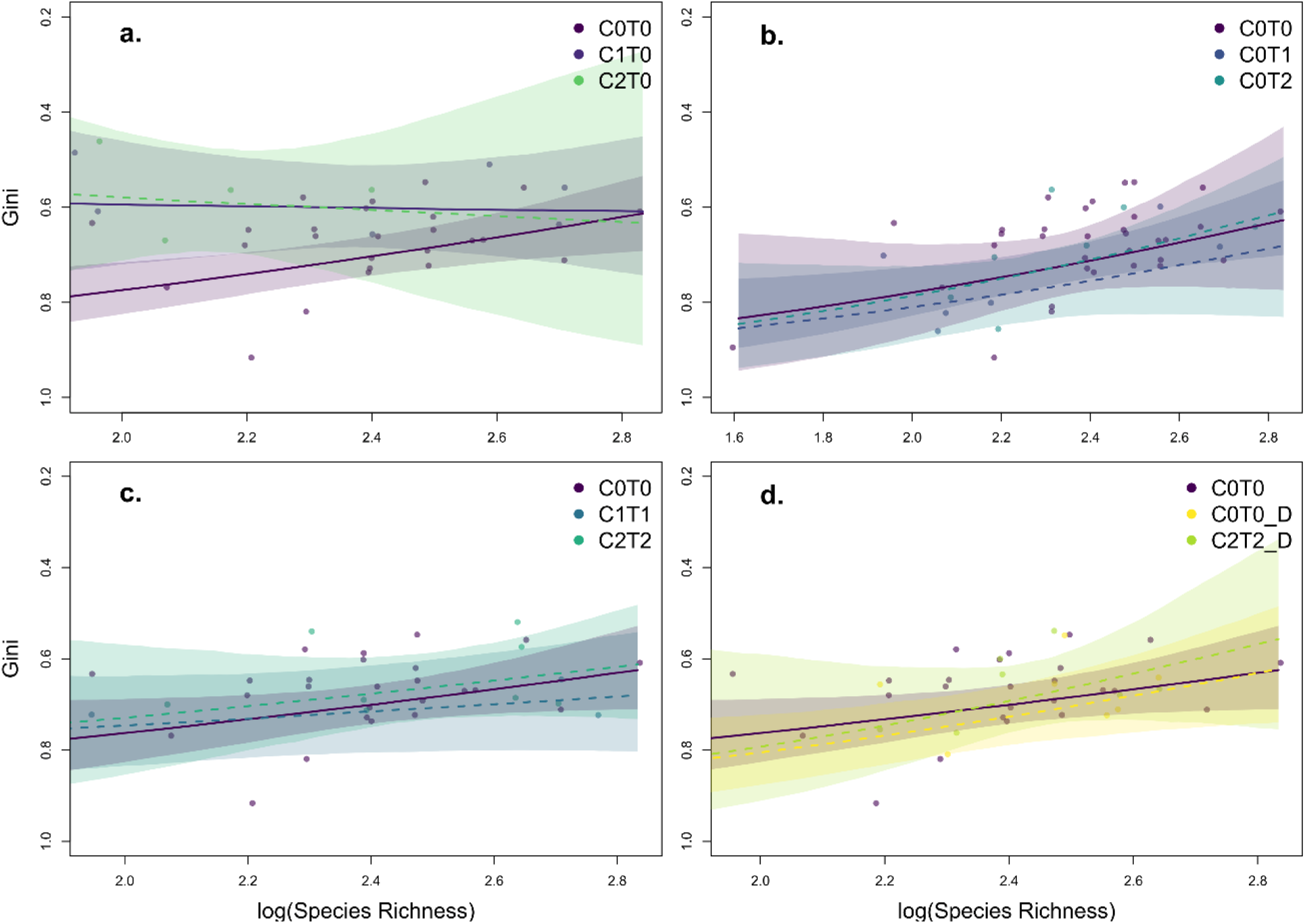
Richness-evenness relationship (RER) responses to climate change treatments after reseeding for control plots vs. **a.** CO_2_ treatments (C1T0: +150 ppm and C2T0: +300 ppm), **b.** temperature treatments (C0T1: +1.5 °C and C0T2: +3°C), **c.** climate change scenarios 1 (C1T1: +150 ppm, + 1.5°C) and 2 (C2T2: +300 ppm and +3°C), and **d.** drought treatments C0T0_D, C2T2_D (implemented only for control and scenario 2 plots). Points, lines, and intervals are as described in the legend to Fig. 2. Data are from vegetation surveys conducted after reseeding (i.e., post-2017), including data only from reseeded plots.

**Table 2:**
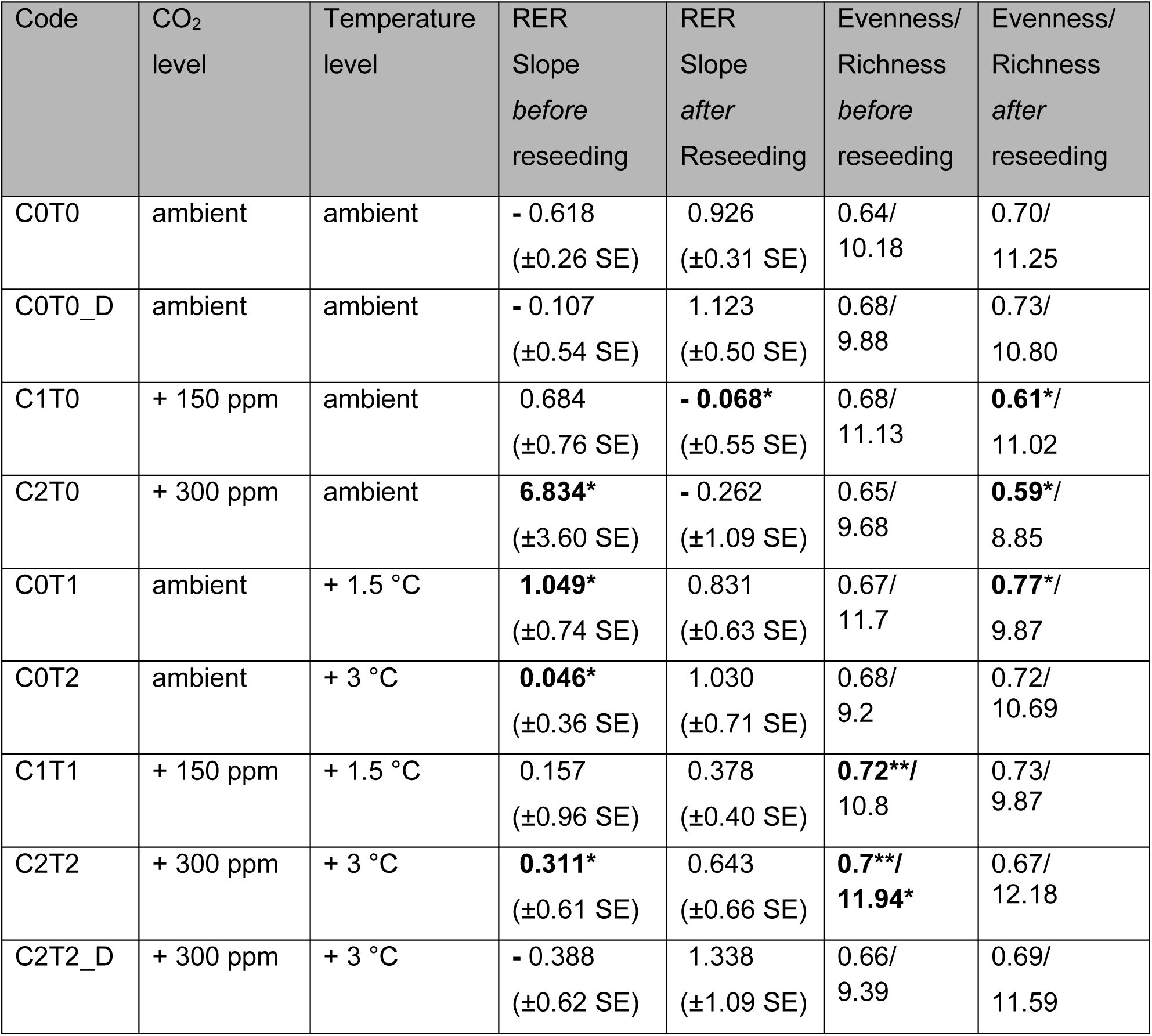
Slopes for the richness evenness relationship (RER) before and after reseeding, values represent model estimates, stars (*) indicate significant difference from control before/after reseeding (C0T0). The significance of differences in RER was identified by comparing the posterior distributions of intercept and slope estimates across the fitted regressions.

| Code | CO<sub>2</sub> level | Temperature level | RER Slope *before* reseeded | RER Slope *after* Reseeding | Evenness/Richness *before* reseeded | Evenness/Richness *after* reseeded |
| --- | --- | --- | --- | --- | --- | --- |
| C0T0 | ambient | ambient | - 0.618<br>(±0.26 SE) | 0.926<br>(±0.31 SE) | 0.64/<br>10.18 | 0.70/<br>11.25 |
| C0T0_D | ambient | ambient | - 0.107<br>(±0.54 SE) | 1.123<br>(±0.50 SE) | 0.68/<br>9.88 | 0.73/<br>10.80 |
| C1T0 | + 150 ppm | ambient | 0.684<br>(±0.76 SE) | - **0.068\***<br>(±0.55 SE) | 0.68/<br>11.13 | **0.61\***/<br>11.02 |
| C2T0 | + 300 ppm | ambient | **6.834\***<br>(±3.60 SE) | - 0.262<br>(±1.09 SE) | 0.65/<br>9.68 | **0.59\***/<br>8.85 |
| C0T1 | ambient | + 1.5 °C | **1.049\***<br>(±0.74 SE) | 0.831<br>(±0.63 SE) | 0.67/<br>11.7 | **0.77\***/<br>9.87 |
| C0T2 | ambient | + 3 °C | **0.046\***<br>(±0.36 SE) | 1.030<br>(±0.71 SE) | 0.68/<br>9.2 | 0.72/<br>10.69 |
| C1T1 | + 150 ppm | + 1.5 °C | 0.157<br>(±0.96 SE) | 0.378<br>(±0.40 SE) | **0.72\*\***/<br>10.8 | 0.73/<br>9.87 |
| C2T2 | + 300 ppm | + 3 °C | **0.311\***<br>(±0.61 SE) | 0.643<br>(±0.66 SE) | **0.7\*\***/<br>**11.94\*** | 0.67/<br>12.18 |
| C2T2_D | + 300 ppm | + 3 °C | - 0.388<br>(±0.62 SE) | 1.338<br>(±1.09 SE) | 0.66/<br>9.39 | 0.69/<br>11.59 |

### Treatments affect richness-evenness relationships

We found significant effects of both temperature and CO_2_ treatments on the relationship between evenness and richness (RER), but interestingly not always in treatments with significant differences in species richness and evenness in comparison to the control (Tab. 2 and Fig. S7-10). Treatment C2T0 (+300 ppm), C0T1 (+1.5°C) and C0T2 (+3°C) showed significant differences in RER slope before reseeding in comparison to the control, while there was no significant difference in evenness or richness alone (Tab. 2). Additionally, both climate scenarios (C1T1 and C2T2), showed no difference in RER slope in comparison to the control (Fig. 3, c), with the only significant difference arising for the intercept (p-value < 0.05) in C2T2 (+300 ppm, + 3 °C), despite significant differences in evenness (C1T1, C2T2) and species richness (C2T2). Combining these scenarios with drought treatments only resulted in small, non-significant differences in RER in comparison to the control (Fig. 2, d).

While the RER in control plots was positive, the relationship became negative in both the +300 ppm and +150 ppm CO_2_ treatments (C1T0, C2T0), with significant differences detected between the +300 ppm (C2T0) vs. control (C0T0) RER slope, as well as between the +300 ppm (C2T0) and the +150 ppm (C1T0) treatment (slope: p-value < 0.05, intercept: p-value < 0.05) (Fig. 3, a and Tab. 2). The temperature treatments (C0T1, C0T2) had similar impacts on RER, though the difference between the control and treatment RER was larger for the +1.5 ° C (C0T1) treatment (slope: p-value < 0.05, intercept: p-value < 0.05) than for the +3°C (C0T2) treatment (intercept: p-value < 0.05), where only the intercept was significantly different from the control (Fig. 2, b). Combining these scenarios with drought treatments only resulted in small, non-significant differences in the RER compared to the control (Fig. 3, d and Tab. 2).

### Reseeding removes differences between treatments

To test whether reseeding can erase, or at least partially alleviate, the effects of climate change on the RER in our system, we also analyzed the RER in all climate change treatments after reseeding. Reseeding took place in early April 2018, 2020, and 2021 – thus, post-reseeding vegetation surveys are available from 2019 and 2022. There is no clear temporal pattern in the evenness and species richness data, despite the increase in evenness after reseeding in 2018 (Fig. 1, c and d).

Additionally, we added year as a categorical fixed effect in our brms models for the RER to control for temporal variation. Recall that the seed mixture used for reseeding was identical to the initial seed mixture; therefore, no new species were actively introduced.

After reseeding, the RER was positive across almost all control and treatment plots, except for the slight negative RER trends in the CO_2_ treatments (Fig. 4a- d). In general, there is only one significant difference between the slope after reseeding between the control and the C1T0 (+150 ppm) treatment, which was significantly lower than in the control plots (Tab. 2).

To investigate the differences in evenness and richness separately before and after reseeding, we compared Control (C0T0) and treatment plots. Evenness, but not richness, was significantly higher in the CO_2_ treatments (C1T0, C2T0), and significantly lower in the lowest temperature treatment (C0T1: +1.5°C). Interestingly, there again was no significant difference in richness or evenness in the C1T0 treatment (+150 ppm) in comparison to the control, despite the significant difference in RER slope (Tab. 2, Fig. 4).

Interestingly, we found that reseeding had different effects on plots depending on whether they had high or low species richness at the time of reseeding (Fig. 5).

**Figure 5:**
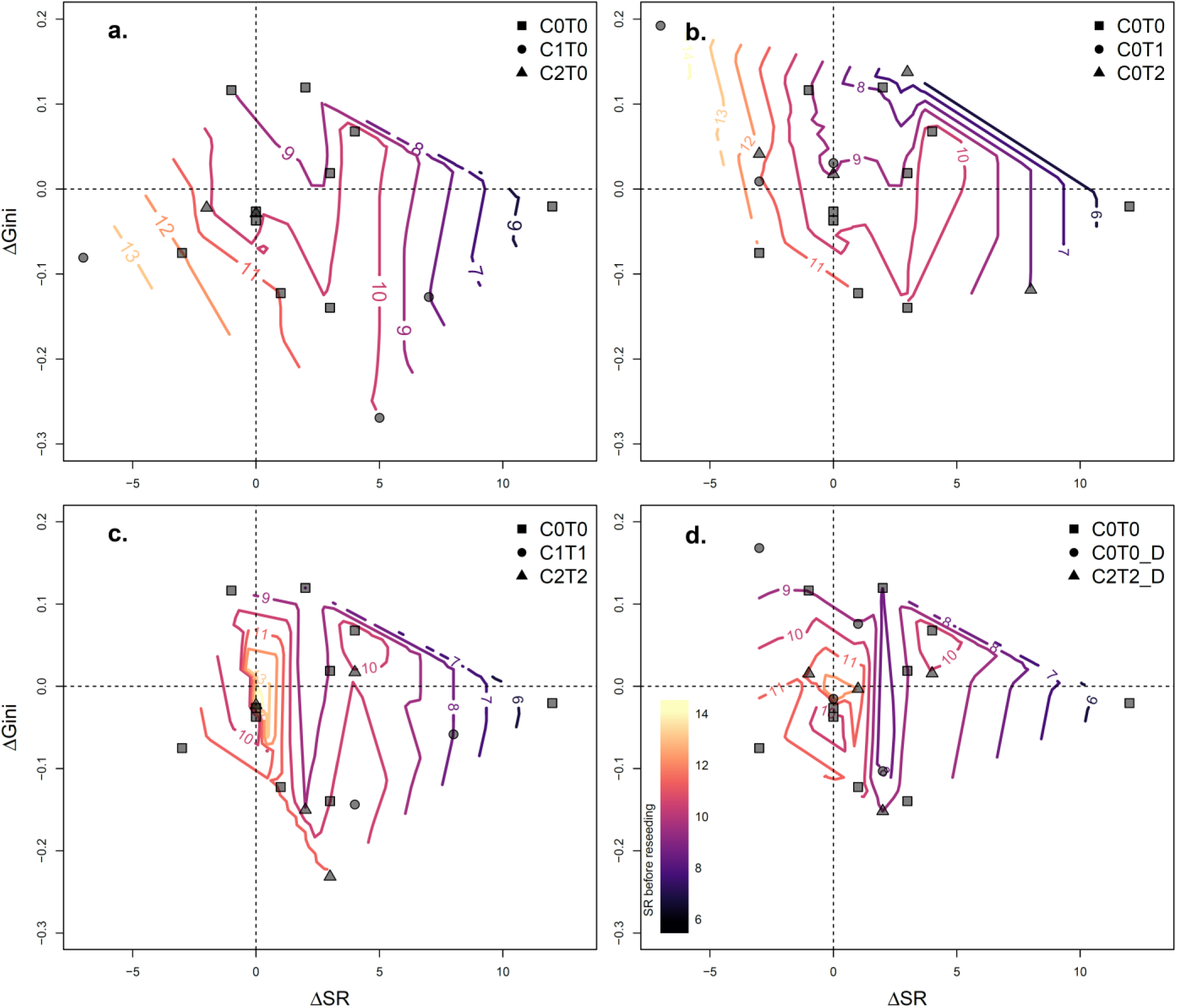
Reseeding enhances richness in low richness plots and reduces richness in high richness plots. Differences (Δ) in species richness and Gini before (≤2017) and after reseeding (≥2019), relative to plot-level species richness observed prior to reseeding. Dashed lines show no difference (Δ = 0) in species richness or evenness before and after reseeding, and colored contour lines show species richness before reseeding. Points show single plots separated by treatment. Shape indicates treatment levels in **a.** CO_2_ treatments (C1T0: +150 ppm and C2T0: +300 ppm), **b.** temperature treatments (C0T1: +1.5 °C and C0T2: +3°C), **c.** climate change scenarios 1 (C1T1: +150 ppm, + 1.5°C) and 2 (C2T2: +300 ppm and +3°C), and **d.** drought treatments C0T0_D, C2T2_D (implemented only for control and scenario 2 plots). Gini is used as an evenness metric, where 0 indicates complete evenness, and 1 indicates complete dominance. Only reseeded plots are included in analyses, with Δ comparisons showing differences in plots from the last survey directly before reseeding, vs. the first survey directly after reseeding.

Reseeding in control plots (C0T0) seemed to reduce evenness. In the CO_2_ treatments (C1T0: +150 ppm, C2T0: +300 ppm), low richness plots increased in richness following reseeding, whereas high richness plots lost species, but gained evenness. In contrast, in temperature treatments (C0T1: +1.5°C, C0T2: +3°C), evenness declined after reseeding in high richness plots, with simultaneous species loss in high richness plots and species gains in low richness plots (Fig. 5, b). The scenarios (C1T1: +150 ppm, +1.5°C; C2T2: +300 ppm, +3°C) gained species following reseeding in high richness plots (C2T2) and in low richness plots (C1T1) (Fig. 5c). In contrast to the other treatments, only 2 plots of the drought treatments (C0T0_D: ambient, + drought, C2T2_D: +300 ppm, +3°C, + drought) lost species following reseeding, while most plots gained species regardless of their species richness prior to reseeding (Fig. 5 d). According to these differences, reseeding of plots seemed to reduce differences in the RER among treatments, but did not restore the negative RER observed in the control plots before reseeding.

### Dominant grass species could cause differences in richness-evenness relationships before and after reseeding

To examine whether the dominant grass species *Arrhenatherum elatius* could be responsible for the differences in the RER before and after reseeding, we looked at how its cover changed as a function of plot-level species richness (Fig. 6). Overall, we found that the cover of *A. elatius* was higher across all treatments after reseeding (Fig. 6, e and f). In the CO_2_ treatments, the cover of *A. elatius* increased in high richness plots and decreased in low richness plots, whereas plot-level richness changed in the opposite direction (Fig. 6, a). The same pattern occurs for evenness and richness before and after reseeding, where high richness plots lost species but gained evenness and low richness plots gained species and decreased in evenness (Fig. 5, a). The temperature treatments (C0T1: +1.5 °C and C0T2: +3 °C) showed similar trends, where high richness plots tended to lose species and gain *A. elatius* cover (Fig. 6, b). In contrast, in the climate change scenarios (C1T1 and C2T2), all plots gained *A. elatius* cover, while also gaining richness, regardless of the richness before reseeding (Fig. 6, c). In the drought treatments (C0T0_D and C2T2_D), 2 plots lost species while gaining *A. elatius* cover (Fig. 6, d).

**Figure 6:**
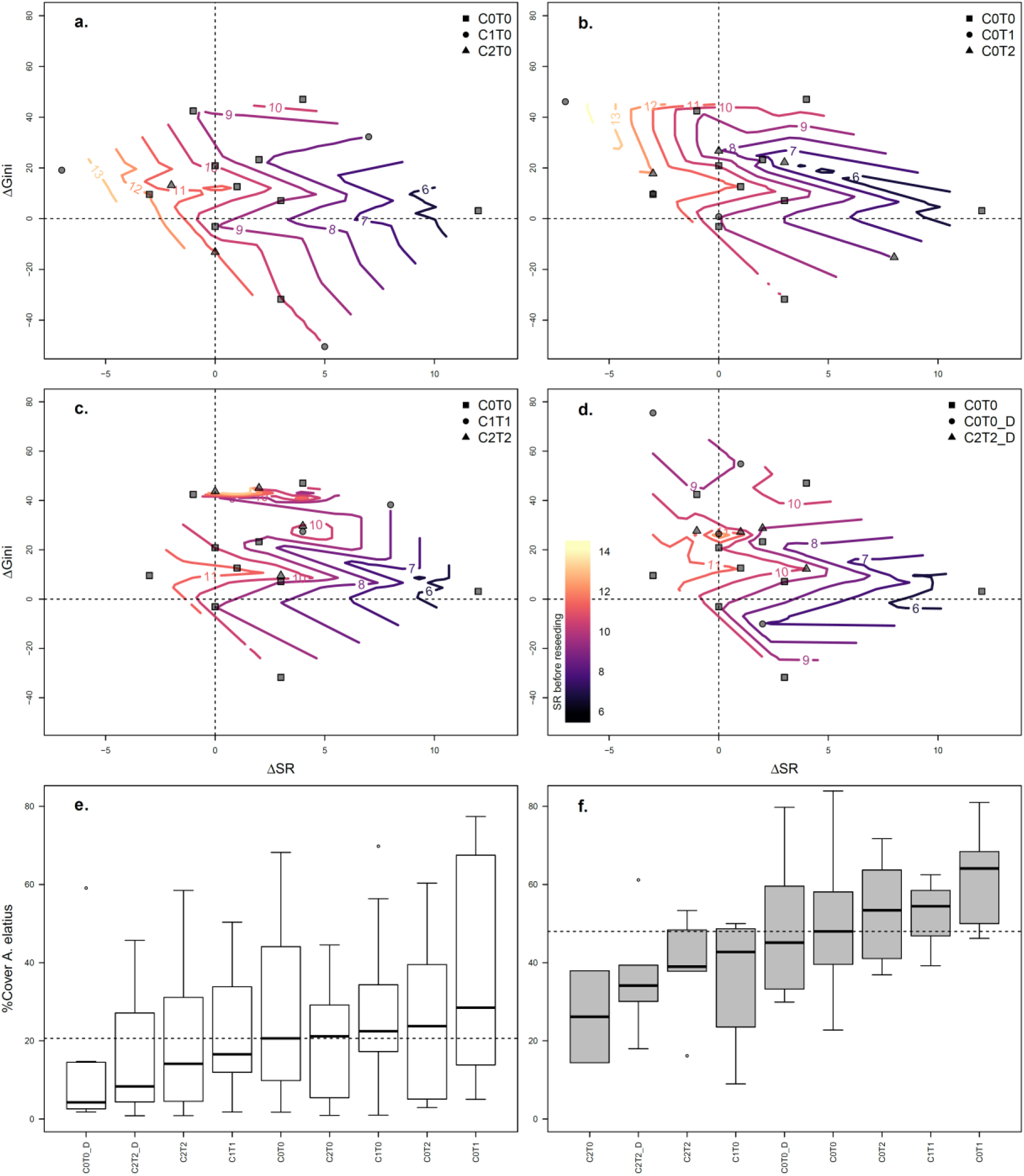
Reseeding enhances *A. elatius* %cover. Differences (Δ) in species richness and *A. elatius* %cover before (≤2017) and after reseeding (≥2019), relative to plot-level species richness observed prior to reseeding. Dashed lines show no change in species richness or *A. elatius* cover, and colored contour lines show species richness before reseeding. Gini is used as an evenness metric, where 0 indicates complete evenness. Points show single plots separated by treatment. Shape indicates treatment levels in **a.** CO_2_ treatments (C1T0: +150 ppm and C2T0: +300 ppm), **b.** temperature treatments (C0T1: +1.5 °C and C0T2: +3°C), **c.** climate change scenarios 1 (C1T1: +150 ppm, + 1.5°C) and 2 (C2T2: +300 ppm and +3°C), and **d.** drought treatments C0T0_D, C2T2_D (implemented only for control and scenario 2 plots). **e., f.** show %cover of *A. elatius* before and after reseeding in all treatments. The dashed line indicates the median of *A. elatius* cover in control plots. Black bars indicate median, boxes 25% and 75% quantiles. Only reseeded plots are included in analyses, with Δ comparisons showing changes in plots from directly before vs. directly after reseeding.

## Discussion

Our results show that, in contrast to evenness or richness alone, richness evenness relationships (RERs) are sensitive to community changes in our experiment, as they contain additional information to means or similar approaches of evenness and richness alone. Also, RERs could indicate changes that differ between high and low richness communities. The RER captures changes in the relationship between community diversity and the distribution of relative abundances across the community. Consequently, the RER may be a first sign of changes in communities affected by climate change, as suggested for evenness (Chapin et al. 2000). Plant communities are expected to have negative RERs (Soininen et al., 2012; Zhang et al., 2012; Hordijk et al., 2023), which corresponds to our results in the non-reseeded control. We show that the RER in our experiment differs significantly between treatments and control before reseeding, even without any corresponding differences in richness or evenness. Therefore, RER represents a signal of community change that differs from what can be gleaned from richness and evenness separately.

This increased sensitivity is especially important in alpine and subalpine regions, as there can be a lag in species responses to climate change, likely longer than in other ecosystems (Alexander et al. 2018; Rumpf et al. 2019). This potential lag suggests that future changes in affected communities could be substantially larger than those observed thus far. There is evidence that the signs of the RER change with environmental context, trophic level, or taxa. Also, Stirling and Wilsey (2001) showed that observed relationships between richness and evenness differed from combinations of evenness and richness, generated by a neutral model. This indicates that biological processes shape the relationship between richness and evenness.

The difference in RER was strongest between temperature treatments and the control, which corresponds to previously reported changes in community composition reported from lowland (Kardol et al.,2010) and alpine grassland ecosystems (Liu et al., 2018). In contrast to our expectations, richness, and evenness did not change significantly, but the slope of the RER in both temperature treatments changed significantly, and even the direction of the RER is positive, in contrast to the negative RER in the non-reseeded control. Other studies from the European Alps found contrasting effects, where increased temperatures only affected plants when treatments were accompanied by drought (Boeck et al., 2016). Like other climate change studies, we found decreased evenness in global change treatments (Kardol et al., 2010), but only in both scenarios, i.e., combinations of increased CO_2_ and temperature levels. Also, we found almost no changes in species richness, other than a significant increase in the climate scenario with +300 ppm CO_2_ and +3°C in temperature. This contrasts with our expectations, where we expected species richness or evenness separately to differ in treatments and controls. The effect of elevated CO_2_ and temperature seemed to change the RER in similar ways in comparison to the control, but their combined effects led to a less pronounced relationship between richness and evenness, with only Scenario 2 (+300ppm, 3°C) being significantly different from the control. This result could indicate that CO_2_ and temperature interact and at least partly counteract one another’s effects on the RER, but not enough to restore the negative RER found in the control. Indeed, some previous studies have found evidence that elevated CO_2_ concentrations can ameliorate the effects of temperature increases (Reyes-Fox et al., 2014; Roy et al., 2016) – potentially because higher CO_2_ concentrations lead to reduced evapotranspiration, thereby buffering the effects of increased temperature (Slawitsch et al., 2020).

The lack of a significant impact of drought treatments on the RER is surprising, and contrasts with some results from other climate change experiments (e.g., White et al., 2014), as well as with our hypotheses. A potential explanation, which matches findings from other managed European grasslands, is that water availability was sufficiently high to prevent drought stress, even within the drought treatments (Hopkins and Del Prado 2007). Indeed, average annual precipitation at our site is particularly high (at 1077 mm/year), and has been increasing with time (see Supplement, Fig. S1). Additionally, recall that in our experiment, drought treatments were implemented as a single, severe event per year, after which re-wetting was applied, as is predicted by climate model forecasts, which predict increased net precipitation with more heavy rain events and dry periods (Gobiet et al., 2014). The short “pulse” may have reduced the effects of drought on species richness and evenness relative to those from longer-term treatments in other experiments – e.g., in the experiment conducted by Kardol et al. (2010), where water content was controlled throughout the duration of the experiment, and the authors found a significant decrease in evenness. Indeed, responses to climate change, especially to drought, seem to differ by local climate and ecosystem properties (Isbell et al., 2015; Boeck et al., 2018).

Another important result is that high and low richness communities often seem to react in opposite directions to reseeding, a community change that is visible through the richness evenness relationship (RER). This result is probably due to *Arrhenaterum elatius*, which was the dominant grass species in all plots after reseeding, and in most plots even before reseeding. In our experiment, reseeding can be thought of as an active restoration method, which is practiced by farmers already today. Reseeding affected the dominance of *A. elatius*, and therefore seemed to change the direction of the RER. The lowest dominance of *A. elatius* seems to indicate no relationship between evenness and richness, or even a slightly positive RER, while an increase in dominance leads to more negative RERs. All treatments in which *A. eliatus* had high dominance, both before and after reseeding, showed a positive RER, i.e., low richness plots with low evenness and high richness plots with high evenness. Therefore, high dominance of A. elatius prior to reseeding was generally associated with low richness and low evenness in plots, both of which increased after reseeding. In contrast, plots that had high richness prior to reseeding also tended to have low evenness and low dominance of *A. elatius* – but, after reseeding, dominance of *A. elatius* generally increased, while richness and evenness both decreased. In sum, reseeding partially alleviated the differences in the RER between treatments and controls, but failed to fully restore the negative RER found in the controls, as suggested in Fig. 1, e. The only exceptions to this pattern were both CO_2_ treatments, where the RER is negative after reseeding, potentially because fewer plots were re-seeded in these treatments (C1T0, +150 ppm: n = 3; C2T0, +300 ppm: n = 2). This difference also shows a major problem in assessing the reseeding treatments in our study. Reseeding mainly took place after the occurrence of “pests” (e.g., ants) and was not part of the experimental design.

Therefore, the number of reseeded plots differs somewhat among treatments, which can make direct comparisons difficult. Recall again that to avoid potential biases in our main analyses, we always limit comparisons either to plots that have not yet been reseeded (i.e., survey years prior to 2017) or to subsets of data for which all plots were reseeded (i.e., survey years after 2017).

Our results from reseeding are in line with recent evidence from animal populations, where captive release is suggested to increase interspecific competition, thereby destabilizing communities (Terui et al. 2023). The results suggest that the large amount of *A. elatius* in the seed mixture used for reseeding (14.9 %, see Table S4) contributes to decreases in plot-level richness and decreases in evenness, especially in high richness communities. This underlines the concerns raised about the ecological risks of intentional release, relocation, or reseeding programs (Araki et al. 2007; Krkosek et al. 2007; Kitada 2014). While most of this evidence is derived from fish, massive releases of plant species also occur in many restoration projects (Laikre et al. 2006) – and the risks that these efforts pose for community diversity, stability, and productivity remain poorly understood. While in our experiment, re- seeding allows low richness communities to gain species, it also appears to drive species loss in high richness communities.

The consequences of different RERs are less clear, but this relationship is probably meaningful, as evenness, like richness, is an important contributor to many different ecosystem properties (Kirwan et al. 2007; Hillebrand et al. 2008; Wittebolle et al. 2009; Wilsey and Potvin 2000). Also, a previous study has found that RERs seem to mediate the relationship between forest productivity and richness (Hordijk et al., 2023), where in low richness systems, uneven communities are more productive than even ones. In contrast, in high richness systems, even communities are more productive than uneven ones. This underpins the importance of our findings of positive RERs as a result of reseeding or climate, which would indicate that the less productive communities (low richness, high evenness and high richness, low evenness) get replaced by more productive communities (low richness, low evenness and high richness, high evenness) (Fig. 1, b). Therefore, changing RERs could greatly affect the productivity of grassland communities. Also, the stability of ecosystems is suggested to be affected not only by evenness and richness but probably depends on the combination of richness and evenness, as they shape different aspects of stability (Hillebrand et al., 2008). As more diverse seed mixtures increase productivity, with corresponding economic benefits in intensively used grasslands (Binder et al. 2018), adjusting evenness accordingly to species richness could further increase these benefits. Taken together, this emphasizes the importance of further investigating the reasons and consequences of different richness evenness relationships (RER).

In summary, we provide evidence that 1) climate change alters the richness- evenness relationship (RER) in our grassland communities, disconnected from changes in evenness and richness, and that 2) reseeding can partially alleviate the changes in the RER across all treatments, but does not restore the negative RER of the control. The RER can reveal insights into community responses to climate change, more sensitive than richness or evenness alone. Like our results from the reseeding show, evenness and species richness in communities is especially important to consider before reseeding, because low and high richness plots can react differently to global change drivers and to management, thereby affecting the direction of the RER, with consequences for productivity and stability (Hillebrand et al., 2008; Hordijk et al., 2023.

As the effects of reseeding differ for each treatment, the RER could help to better match restoration treatments to global change drivers. Our results suggest that incorporating the RER into management decisions might be important and that carefully chosen seed mixtures could ameliorate the impacts of climate change. Our results show that reseeding with the original seed mixture for buffering biodiversity responses to climate change can lead to less productive meadows by supporting the dominant species. Therefore, a more nuanced approach integrating evenness, richness, and their relation in seed mixtures could be an important step forward to buffer the effects of climate change, not only in intensively used grasslands.

## Data Availability

All data and code needed to reproduce the analyses presented here can be found at https://doi.org/10.5281/zenodo.14134001.

## Acknowledgements

We are grateful for the support of the staff at RBG, particularly for their assistance with the vegetation surveys conducted prior to 2022.

## Author contributions

LKM developed the idea for this study. Vegetation surveys in 2022 were conducted by ATC, AK, and LKM. LKM wrote the first draft of the paper. ATC, AK, AS, LG, and LKM contributed to revisions.

## References

1. Alexander, Jake M.; Chalmandrier, Loïc; Lenoir, Jonathan; Burgess, Treena I.; Essl, Franz; Haider, Sylvia et al. (2018): Lags in the response of mountain plant communities to climate change. In: Global Change Biology 24 (2), S. 563–579. DOI: 10.1111/gcb.13976.

2. Araki, Hitoshi; Cooper, Becky; Blouin, Michael S. (2007): Genetic effects of captive breeding cause a rapid, cumulative fitness decline in the wild. In: Science (New York, N.Y.) 318 (5847), S. 100–103. DOI: 10.1126/science.1145621.

3. Binder, Seth; Isbell, Forest; Polasky, Stephen; Catford, Jane A.; Tilman, David (2018): Grassland biodiversity can pay. In: Proceedings of the National Academy of Sciences of the United States of America 115 (15), S. 3876–3881. DOI: 10.1073/pnas.1712874115.

4. Blowes, Shane A.; Daskalova, Gergana N.; Dornelas, Maria; Engel, Thore; Gotelli, Nicholas J.; Magurran, Anne E. et al. (2022): Local biodiversity change reflects interactions among changing abundance, evenness, and richness. In: Ecology, e3820. DOI: 10.1002/ecy.3820.

5. de Boeck, Hans J., Bassin, Seraina; Verlinden, Maya; Zeiter, Michaela; Hiltbrunner, Erika (2016): Simulated heat waves affected alpine grassland only in combination with drought. In: The New phytologist 209 (2), S. 531–541. DOI: 10.1111/nph.13601.

6. de Boeck, Hans J.; Bloor, Juliette M. G.; Kreyling, Juergen; Ransijn, Johannes C. G.; Nijs, Ivan; Jentsch, Anke; Zeiter, Michaela (2018): Patterns and drivers of biodiversity-stability relationships under climate extremes. In: Journal of Ecology 106 (3), S. 890–902. DOI: 10.1111/1365-2745.12897.

7. Bürkner, Paul-Christian (2017): brms : An R Package for Bayesian Multilevel Models Using Stan. In: Journal of Statistical Software 80 (1). DOI: 10.18637/jss.v080.i01.

8. Bütof, Astrid; von Riedmatten, Lars R.; Dormann, Carsten F.; Scherer-Lorenzen, Michael; Welk, Erik; Bruelheide, Helge (2012): The responses of grassland plants to experimentally simulated climate change depend on land use and region. In: Global Change Biology 18 (1), S. 127–137. DOI: 10.1111/j.1365-2486.2011.02539.x.

9. Chao, Anne; Ricotta, Carlo (2019): Quantifying evenness and linking it to diversity, beta diversity, and similarity. In: Ecology 100 (12), e02852. DOI: 10.1002/ecy.2852.

10. Chapin, F. S.; Zavaleta, E. S.; Eviner, V. T.; Naylor, R. L.; Vitousek, P. M.; Reynolds, H. L. et al. (2000): Consequences of changing biodiversity. In: Nature 405 (6783), S. 234–242. DOI: 10.1038/35012241.

11. Clare, David S.; Culhane, Fiona; Robinson, Leonie A. (2022): Secondary production increases with species richness but decreases with species evenness of benthic invertebrates. In: Oikos 2022 (4), e08629. DOI: 10.1111/oik.08629.

12. Deltedesco, Evi; Keiblinger, Katharina M.; Naynar, Maria; Piepho, Hans-Peter; Gorfer, Markus; Herndl, Markus et al. (2019): Trace gas fluxes from managed grassland soil subject to multifactorial climate change manipulation. In: Applied Soil Ecology 137, S. 1–11. DOI: 10.1016/j.apsoil.2018.12.023.

13. Eurostat (2020): Share of Main Land Types in Utilised Agricultural Area (UAA) by NUTS 2 Regions.

14. Fussmann, Katarina E.; Schwarzmüller, Florian; Brose, Ulrich; Jousset, Alexandre; Rall, Björn C. (2014): Ecological stability in response to warming. In: Nature Climate Change 4 (3), S. 206–210. DOI: 10.1038/nclimate2134.

15. Gobiet, Andreas; Kotlarski, Sven; Beniston, Martin; Heinrich, Georg; Rajczak, Jan; Stoffel, Markus (2014): 21st century climate change in the European Alps--a review (493). Online verfügbar unter https://reader.elsevier.com/reader/sd/pii/S0048969713008188?token=2C460EB8248840AF774E2D97D8FFB9F189D91A59161EA357B16069B5150214F5DCC94AC9ACB297E72D33EF6ED245C77E&originRegion=eu-west-1&originCreation=20230227135403.

16. Gruner, Daniel S.; Bracken, Matthew E. S.; Berger, Stella A.; Eriksson, Britas Klemens; Gamfeldt, Lars; Matthiessen, Birte et al. (2017): Effects of experimental warming on biodiversity depend on ecosystem type and local species composition. In: Oikos 126 (1), S. 8–17. DOI: 10.1111/oik.03688.

17. Herndl, Markus; Pötsch, Erich; Bohner, Andreas; Kandolf, Matthias (2011): Lysimeter als Bestandteil eines technischen Versuchskonzeptes zur Simulation der Erderwärmung im Grünland.

18. Hillebrand, Helmut; Bennett, Danuta M.; Cadotte, Marc W. (2008): Consequences of dominance: a review of evenness effects on local and regional ecosystem processes. In: Ecology 89 (6), S. 1510–1520. DOI: 10.1890/07-1053.1.

19. Hopkins, A.; Del Prado, A. (2007): Implications of climate change for grassland in Europe: impacts, adaptations and mitigation options: a review. In: Grass and Forage Science 62 (2), S. 118–126. DOI: 10.1111/j.1365-2494.2007.00575.x.

20. Hordijk, Iris; Maynard, Daniel S.; Hart, Simon P.; Lidong, Mo; Steege, Hans ter; Liang, Jingjing et al. (2023): Evenness mediates the global relationship between forest productivity and richness. In: Journal of Ecology, Artikel 1365-2745.14098. DOI: 10.1111/1365-2745.14098.

21. Isbell, Forest; Craven, Dylan; Connolly, John; Loreau, Michel; Schmid, Bernhard; Beierkuhnlein, Carl et al. (2015): Biodiversity increases the resistance of ecosystem productivity to climate extremes. In: Nature 526 (7574), S. 574–577. DOI: 10.1038/nature15374.

22. Kardol, Paul; Campany, Courtney E.; Souza, Lara; Norby, Richard J.; Weltzin, Jake F.; Classen, Aime T. (2010): Climate change effects on plant biomass alter dominance patterns and community evenness in an experimental old-field ecosystem. In: Global Change Biology 16 (10), S. 2676–2687. DOI: 10.1111/j.1365-2486.2010.02162.x.

23. Kirwan, L.; Lüscher, A.; Sebastià, M. T.; Finn, J. A.; Collins, R. P.; Porqueddu, C. et al. (2007): Evenness drives consistent diversity effects in intensive grassland systems across 28 European sites. In: Journal of Ecology 95 (3), S. 530–539. DOI: 10.1111/j.1365-2745.2007.01225.x.

24. Kitada, Shuichi (2014): Japanese chum salmon stock enhancement: current perspective and future challenges. In: Fisheries Science 80 (2), S. 237–249. DOI: 10.1007/s12562-013-0692- 8.

25. Klein, Julia A.; Harte, John; Zhao, Xin-Quan (2004): Experimental warming causes large and rapid species loss, dampened by simulated grazing, on the Tibetan Plateau. In: Ecology letters 7 (12), S. 1170–1179. DOI: 10.1111/j.1461-0248.2004.00677.x.

26. Krkosek, Martin; Ford, Jennifer S.; Morton, Alexandra; Lele, Subhash; Myers, Ransom A.; Lewis, Mark A. (2007): Declining wild salmon populations in relation to parasites from farm salmon. In: Science (New York, N.Y.) 318 (5857), S. 1772–1775. DOI: 10.1126/science.1148744.

27. Ladouceur, Emma; Isbell, Forest; Clark, Adam T.; Harpole, W. Stanley; Reich, Peter B.; Tilman, G. David; Chase, Jonathan M. (2023): The recovery of plant community composition following passive restoration across spatial scales. In: Journal of Ecology 111 (4), S. 814–829. DOI: 10.1111/1365-2745.14063.

28. Laikre, Linda; Palme, Anna; Josefsson, Melanie; Utter, Fred; Ryman, Nils (2006): Release of alien populations in Sweden. In: Ambio 35 (5), S. 255–261. DOI: 10.1579/05-a-060r.1.

29. Liu, Huiying; Mi, Zhaorong; Lin, Li; Wang, Yonghui; Zhang, Zhenhua; Zhang, Fawei et al. (2018): Shifting plant species composition in response to climate change stabilizes grassland primary production. In: Proceedings of the National Academy of Sciences of the United States of America 115 (16), S. 4051–4056. DOI: 10.1073/pnas.1700299114.

30. Lorenz, M. O. (1905): Methods of Measuring the Concentration of Wealth. In: Publications of the American Statistical Association 9 (70), S. 209. DOI: 10.2307/2276207.

31. Navas, Marie-Laure; Garnier, Eric; Austin, Michael P.; Viaud, Agnès; Gifford, Roger M. (2002): Seeking a sound index of competitive intensity: Application to the study of biomass production under elevated CO 2 along a nitrogen gradient. In Austral Ecology 27 (4), pp. 463–473. DOI: 10.1046/j.1442-9993.2002.01201.x.

32. Niklaus, P. A.; Leadley, P. W.; Schmid, B.; Körner, Ch. (2001): A LONG-TERM FIELD STUDY ON BIODIVERSITY × ELEVATED CO 2 INTERACTIONS IN GRASSLAND. In Ecological Monographs 71 (3), pp. 341–356. DOI: 10.1890/0012-9615(2001)071[0341:ALTFSO]2.0.CO;2.

33. Peratoner, Giovanni; Pötsch, Erich M. (2019): Methods to describe the botanical composition of vegetation in grassland research. In: Die Bodenkultur: Journal of Land Management, Food and Environment 70 (1), S. 1–18. DOI: 10.2478/boku-2019-0001.

34. Piepho, H.-P.; Herndl, M.; Pötsch, E. M.; Bahn, M. (2017): Designing an experiment with quantitative treatment factors to study the effects of climate change. In: Journal of Agronomy and Crop Science 203 (6), S. 584–592. DOI: 10.1111/jac.12225.

35. Potvin, Catherine; Vasseur, Liette (1997): LONG-TERM CO 2 ENRICHMENT OF A PASTURE COMMUNITY: SPECIES RICHNESS, DOMINANCE, AND SUCCESSION. In Ecology 78 (3), pp. 666–677. DOI: 10.1890/0012-9658(1997)078[0666:LTCEOA]2.0.CO;2.

36. Potvin, Catherine; Chapin, F. Stuart; Gonzalez, Andrew; Leadley, Paul; Reich, Peter; Roy, Jacques (2007): Plant Biodiversity and Responses to Elevated Carbon Dioxide. In Josep G. Canadell (Ed.): Terrestrial Ecosystems in a Changing World. With assistance of Diane E. Pataki, Louis F. Pitelka. Berlin, Heidelberg: Springer (Global Change - the IGBP Ser), pp. 103–112.

37. Pötsch, Erich; Herndl, Markus; Bahn, Michael; Schaumberger, Andreas; Schweiger, Medardus; Kandolf, Matthias et al. (2020): ClimGrass -ein innovatives Freilandexperiment zur Erforschung der Folgen des Klimawandels im Grünland.

38. Ramsier, Dieter; Connolly, John; Bazzaz, Fakhri A. (2005): Carbon dioxide regime, species identity and influence of species initial abundance as determinants of change in stand biomass composition in five-species communities: an investigation using a simplex design and RGRD analysis. In J Ecology 93 (3), pp. 502–511. DOI: 10.1111/j.1365-2745.2005.00999.x.

39. Reich, Peter B. (2009): Elevated CO_2_ reduces losses of plant diversity caused by nitrogen deposition. In Science (New York, N.Y.) 326 (5958), pp. 1399–1402. DOI: 10.1126/science.1178820.

40. Reyes-Fox, Melissa; Steltzer, Heidi; Trlica, M. J.; McMaster, Gregory S.; Andales, Allan A.; LeCain, Dan R.; Morgan, Jack A. (2014): Elevated CO_2_ further lengthens growing season under warming conditions. In: Nature 510 (7504), S. 259–262. DOI: 10.1038/nature13207.

41. Roy, Jacques; Picon-Cochard, Catherine; Augusti, Angela; Benot, Marie-Lise; Thiery, Lionel; Darsonville, Olivier et al. (2016): Elevated CO_2_ maintains grassland net carbon uptake under a future heat and drought extreme. In: Proceedings of the National Academy of Sciences 113 (22), S. 6224–6229. DOI: 10.1073/pnas.1524527113.

42. Rumpf, Sabine B.; Hülber, Karl; Wessely, Johannes; Willner, Wolfgang; Moser, Dietmar; Gattringer, Andreas et al. (2019): Extinction debts and colonization credits of non-forest plants in the European Alps. In: Nature communications 10 (1), S. 4293. DOI: 10.1038/s41467-019-12343-x.

43. Ryals, Rebecca; Hartman, Melannie D.; Parton, William J.; DeLonge, Marcia S.; Silver, Whendee L. (2015): Long-term climate change mitigation potential with organic matter management on grasslands. In: Ecological Applications 25 (2), S. 531–545. DOI: 10.1890/13-2126.1.

44. Samson, Fred; Knopf, Fritz (1994): Prairie Conservation in North America. In: BioScience 44 (6), S. 418–421. DOI: 10.2307/1312365.

45. Slodowicz, Daniel; Durbecq, Aure; Ladouceur, Emma; Eschen, René; Humbert, Jean-Yves; Arlettaz, Raphaël (2023): The relative effectiveness of different grassland restoration methods: A systematic literature search and meta-analysis. In: Ecological Solutions and Evidence 4 (2), Artikel e12221. DOI: 10.1002/2688-8319.12221.

46. Smith, Benjamin; Wilson, Bastow J. (1996): A Consumer’s Guide to Evenness Indices. In: Oikos May (Vol. 76), pp. 70–82.

47. Soininen, Janne; Passy, Sophia; Hillebrand, Helmut (2012): The relationship between species richness and evenness: a meta-analysis of studies across aquatic ecosystems. In: Oecologia 169 (3), S. 803–809. DOI: 10.1007/s00442-011-2236-1.

48. Stevens, Nicola; Bond, William; Feurdean, Angelica; Lehmann, Caroline E.R. (2022): Grassy Ecosystems in the Anthropocene. In: Annual Review of Environment and Resources 47 (1), S. 261–289. DOI: 10.1146/annurev-environ-112420-015211.

49. Stirling, Gray; Wilsey, Brian (2001): Empirical Relationships between Species richness, Evenness, and Proportional Diversity. In: The American naturalist 158, S. 286–299. DOI: 10.1086/321317.

50. Tuomisto, Hanna (2012): An updated consumer’s guide to evenness and related indices. In Oikos 121 (8), pp. 1203–1218. DOI: 10.1111/j.1600-0706.2011.19897.x.

51. Terui, Akira; Urabe, Hirokazu; Senzaki, Masayuki; Nishizawa, Bungo (2023): Intentional release of native species undermines ecological stability. In: Proceedings of the National Academy of Sciences of the United States of America 120 (7), e2218044120. DOI: 10.1073/pnas.2218044120.

52. van Sundert, Kevin; Arfin Khan, Mohammed A. S.; Bharath, Siddharth; Buckley, Yvonne M.; Caldeira, Maria C.; Donohue, Ian et al. (2021): Fertilized graminoids intensify negative drought effects on grassland productivity. In: Global Change Biology 27 (11), S. 2441–2457. DOI: 10.1111/gcb.15583.

53. Veronika Slawitsch; Steffen Birk; Markus Herndl; Erich M Pötsch (Hg.) (2019): Einfluss des Klimawandels auf die Bodenwasserbilanz im inneralpinen Grünland. 21. Alpenländisches Expertenforum.

54. WallisDeVries, Michiel F.; Poschlod, Peter; Willems, Jo H. (2002): Challenges for the conservation of calcareous grasslands in northwestern Europe: integrating the requirements of flora and fauna. In: Biological Conservation 104, S. 265–273.

55. Wang, Hao; Liu, Huiying; Cao, Guangmin; Ma, Zhiyuan; Li, Yikang; Zhang, Fawei et al. (2020): Alpine grassland plants grow earlier and faster but biomass remains unchanged over 35 years of climate change. In: Ecology letters 23 (4), S. 701–710. DOI: 10.1111/ele.13474.

56. Wang, Xiao-Yan; Ge, Yuan; Gao, Song; Chen, Tong; Wang, Jiang; Yu, Fei-Hai (2021): Evenness alters the positive effect of species richness on community drought resistance via changing complementarity. In: Ecological Indicators 133, S. 108464. DOI: 10.1016/j.ecolind.2021.108464.

57. White, Shannon R.; Bork, Edward W.; Cahill, James F. (2014): Direct and indirect drivers of plant diversity responses to climate and clipping across northern temperate grassland. In: Ecology 95 (11), S. 3093–3103. DOI: 10.1890/14-0144.1.

58. Wilsey, Brian J.; Potvin, Catherine (2000): Biodiversity and ecosystem functioning: Importance of species evenness in an old field. In: Ecology 81 (4), S. 887–892. DOI: 10.1890/0012-9658(2000)081[0887:BAEFIO]2.0.CO;2.

59. Wittebolle, Lieven; Marzorati, Massimo; Clement, Lieven; Balloi, Annalisa; Daffonchio, Daniele; Heylen, Kim et al. (2009): Initial community evenness favours functionality under selective stress. In: Nature 458 (7238), S. 623–626. DOI: 10.1038/nature07840.

60. Zavaleta, Erika S.; Shaw, M. Rebecca; Chiariello, Nona R.; Mooney, Harold A.; Field, Christopher B. (2003): Additive effects of simulated climate changes, elevated CO_2_, and nitrogen deposition on grassland diversity. In: Proceedings of the National Academy of Sciences of the United States of America 100 (13), S. 7650–7654. DOI: 10.1073/pnas.0932734100.

61. Zhang, Hui; John, Robert; Peng, Zechen; Yuan, Jianli; Chu, Chengjin; Du, Guozhen; Zhou, Shurong (2012): The relationship between species richness and evenness in plant communities along a successional gradient: a study from sub-alpine meadows of the Eastern Qinghai-Tibetan Plateau, China. In: PloS one 7 (11), e49024. DOI: 10.1371/journal.pone.0049024.

