## Supplement for "Climate change and reseeding shape richness-evenness relationships in a subalpine grassland experiment"

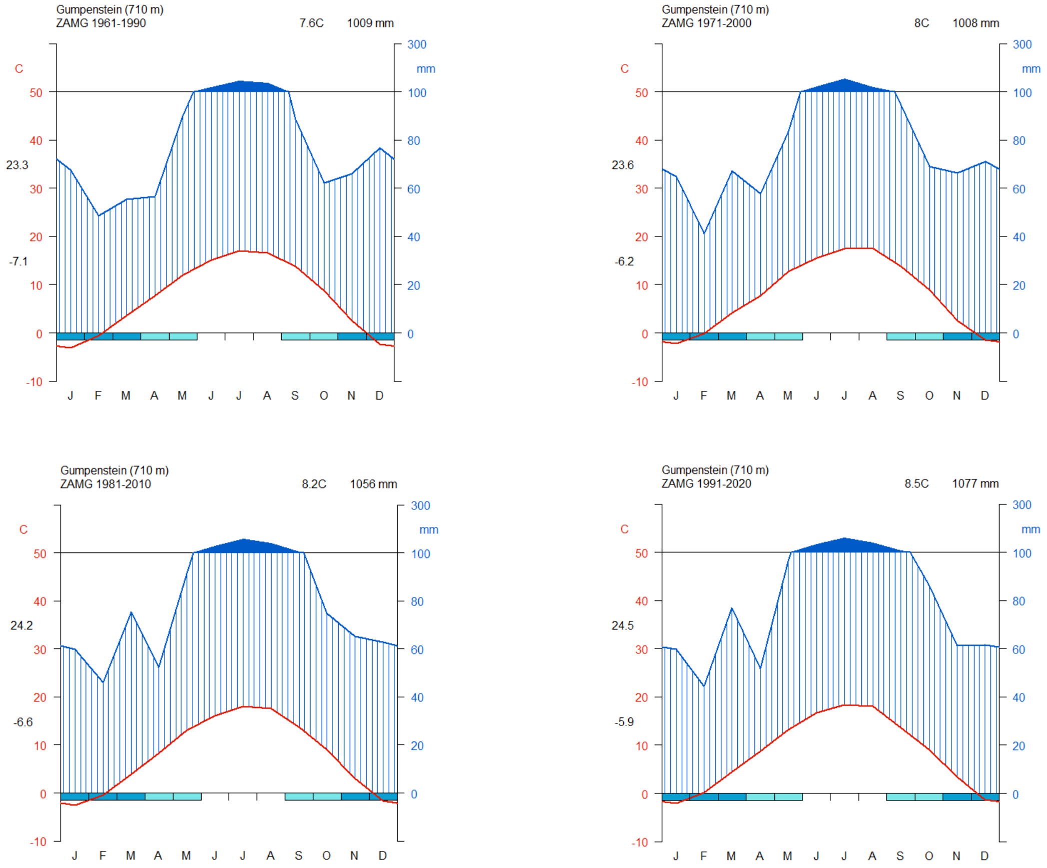

**Figure S1: Climate diagrams for 30 year-periods from 1961 to 2020 for the location of the ClimGrass experiment (Raumberg)**

**Table S1: p-values RER CO2 levels, derived from brms posterior distributions**

| Comparison | type | p-value | |
|---|---|---|---|
| 300_0_No; 0_0_No | slope | 0.021 | * |
| 150_0_No; 0_0_No | slope | 0.065 | |
| 300_0_No; 150_0_No | slope | 0.046 | * |
| 300_0_No; 0_0_No | intercept | 0.02 | * |
| 150_0_No; 0_0_No | intercept | 0.06 | |
| 300_0_No; 150_0_No | intercept | 0.047 | * |
| 300_1_No; 0_1_No | slope | 0.143 | |
| 150_1_No; 0_1_No | slope | 0.065 | |

| | | | |
|---|---|---|---|
| **300_1_No; 150_1_No** | **slope** | 0.411 | |
| **300_1_No; 0_1_No** | **intercept** | 0.102 | |
| **150_1_No; 0_1_No** | **intercept** | 0.035 | * |
| **300_0_No; 150_1_No** | **intercept** | 0.407 | |

**Table S2: p-values for Tmp levels, derived from brms posterior distributions**

| **Comparison** | **type** | **p-value** | |
|---|---|---|---|
| **3_0_No; 0_0_No** | **slope** | 0.05025 | |
| **1.5_0_No; 0_0_No** | **slope** | 0.0145 | * |
| **3_0_No; 1.5_0_No** | **slope** | 0.106 | |
| **3_0_No; 0_0_No** | **intercept** | 0.036 | * |
| **1.5_0_No; 0_0_No** | **intercept** | 0.01375 | * |
| **3_0_No; 1.5_0_No** | **intercept** | 0.10125 | |
| **3_1_No; 0_1_No** | **slope** | 0.43075 | |
| **1.5_1_No; 0_1_No** | **slope** | 0.46425 | |
| **3_1_No; 1.5_1_No** | **slope** | 0.4145 | |
| **3_1_No; 0_1_No** | **intercept** | 0.4325 | |
| **1.5_1_No; 0_1_No** | **intercept** | 0.486 | |
| **3_1_No; 1.5_1_No** | **intercept** | 0.456 | |

**Table S3: p-values Scenarios, derived from brms posterior distributions**

| **Comparison** | **type** | **p-value** | |
|---|---|---|---|
| **2_0_No; 0_0_No** | **slope** | 0.05375 | |
| **1_0_No; 0_0_No** | **slope** | 0.1845 | |
| **2_0_No; 1_0_No** | **slope** | 0.56625 | |
| **2_0_No; 0_0_No** | **intercept** | 0.04075 | * |
| **1_0_No; 0_0_No** | **intercept** | 0.14425 | |

| | | |
|---|---|---|
| **2_0_No; 1_0_No** | **intercept** | 0.4445 |
| **2_1_No; 0_1_No** | **slope** | 0.423 |
| **1_1_No; 0_1_No** | **slope** | 0.232 |
| **2_1_No; 1_1_No** | **slope** | 0.359 |
| **2_1_No; 0_1_No** | **intercept** | 0.40325 |
| **1_1_No; 0_1_No** | **intercept** | 0.244 |
| **2_1_No; 1_1_No** | **intercept** | 0.39775 |
| **0_0_No; 0_0_Yes** | **slope** | 0.149 |
| **0_0_No; 2_0_Yes** | **slope** | 0.309 |
| **0_0_Yes; 2_0_Yes** | **slope** | 0.36325 |
| **0_0_No; 0_0_Yes** | **intercept** | 0.1175 |
| **0_0_No; 2_0_Yes** | **intercept** | 0.3015 |
| **0_0_Yes; 2_0_Yes** | **intercept** | 0.34075 |
| **0_1_No; 0_1_Yes** | **slope** | 0.315 |
| **0_1_No; 2_1_Yes** | **slope** | 0.3125 |
| **0_1_Yes; 2_1_Yes** | **slope** | 0.42325 |
| **0_1_No; 0_1_Yes** | **intercept** | 0.28125 |
| **0_1_No; 2_1_Yes** | **intercept** | 0.32075 |
| **0_1_Yes; 2_1_Yes** | **intercept** | 0.4515 |

13

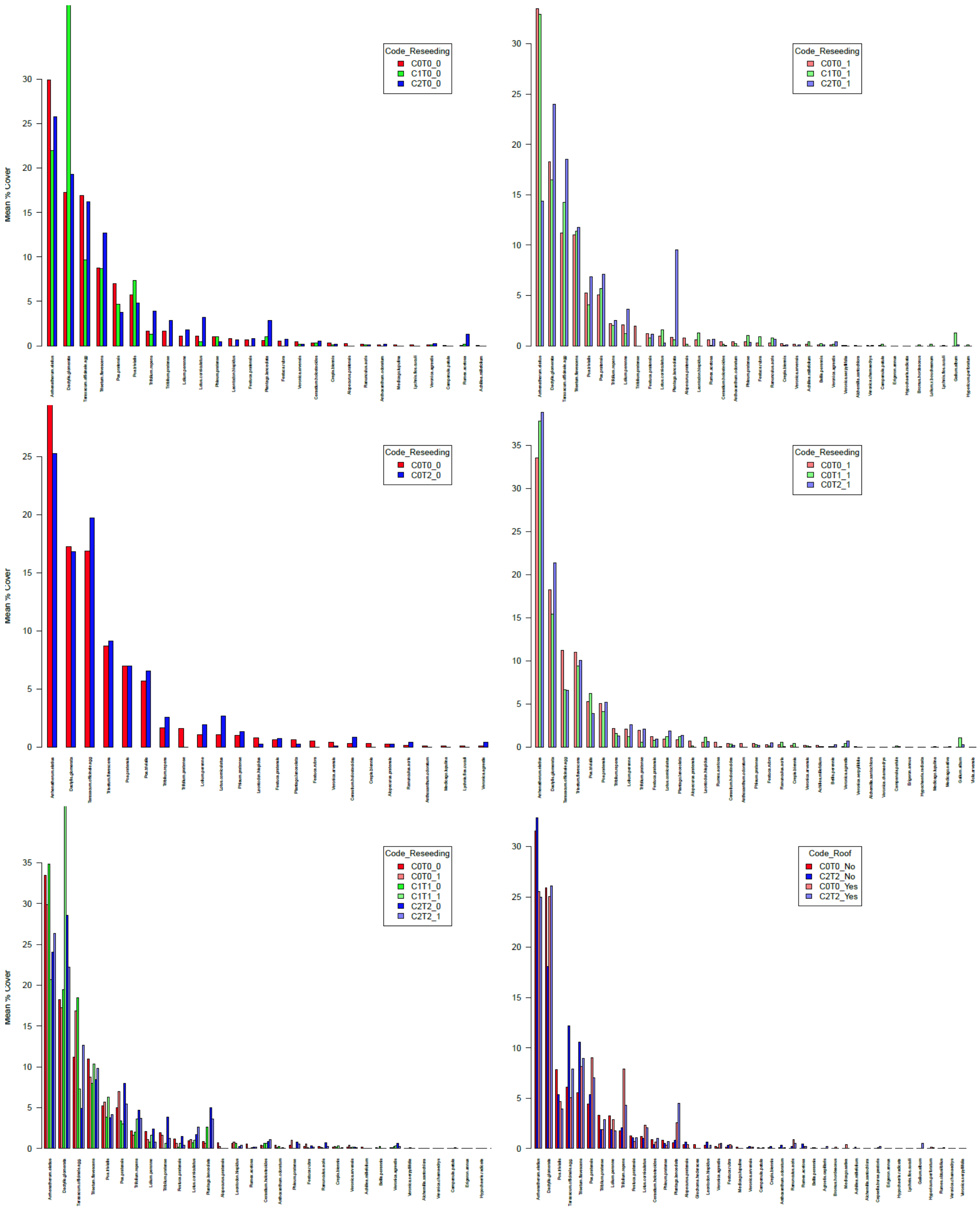

**Figure S2:** Mean % cover for each species in **a.** $CO^2$ treatments (+150 ppm and +300 ppm), **b.** temperature (+1.5 °C and +3°C), **c.** scenario 1 (+150 ppm, + 1.5°C) and scenario 2 (+300 ppm and +3°C), and **d.** drought treatments in control and scenario 2. Abundance distributions for each treatment were calculated with the mean cover per species without cover normalization.

**Table S4: Seed mixture ("DW B")** "ÖAG Dauerwiesenmischung B" A mixture for up to 3 cuts per year, for well water-supplied, moderate to deep meadow sites in the alpine foreland, in valleys and basins and in climatically favorable locations up to an altitude of 800 m.

| Species | % |
|---|---|
| *Lolium perenne* | 9.3 |
| *Arrhenatherum elatius* | 14.9 |
| *Trisetum flavescens* | 3.7 |
| *Dactylis glomerata* | 9 |
| *Festuca rubra* | 5.6 |
| *Phleum pratense* | 7.5 |
| *Alopecurus pratensis* | 5.6 |
| *Poa pratensis* | 20.2 |
| *Festuca pratensis* | 11.2 |
| *Lotus corniculatus* | 5.6 |
| *Trifolium repens* | 7.5 |

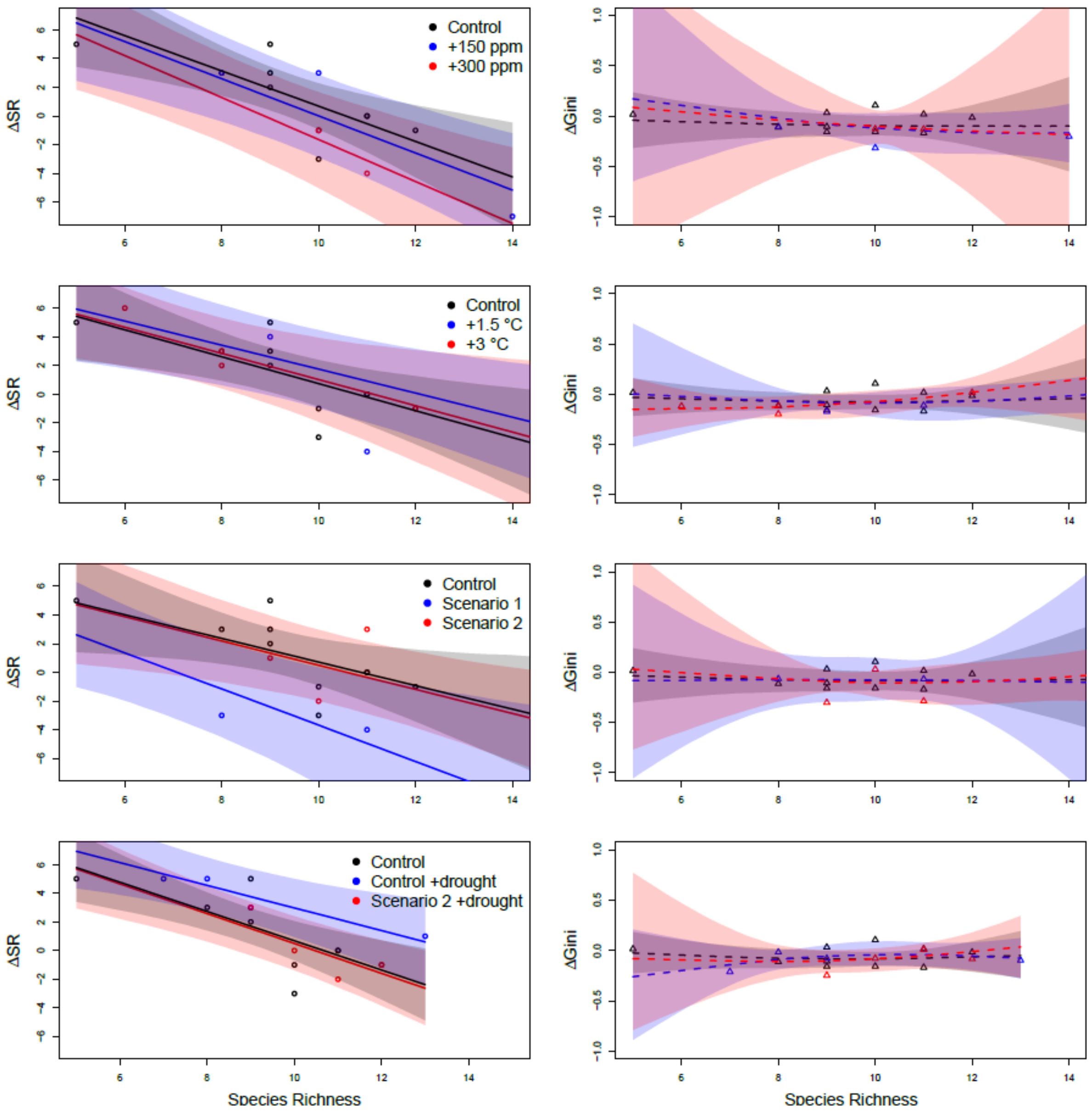

**Figure S4: brms models of the difference in species richness (left panel) and Gini (right panel), before (2017) and after (2019) reseeding.** Regressions and confidence intervals are shown. Color indicates treatment levels in **a.** $CO^2$ treatments (+150 ppm and +300 ppm), **b.** temperature treatments (+1.5 °C and +3°C), **c.** scenario 1 (+150 ppm, + 1.5°C) and scenario 2 (+300 ppm and +3°C), and **d.** drought treatments (in control and scenario 2). P-values of difference between treatments can be found in **Tab. S5 and S6.**

**Table S5: p-values for differences in slopes of ∆species richness and ∆Gini (before and after reseeding) vs. species richness before reseeding, derived from brms posterior distributions**

| **Comparison** | **type** | **p-value** |
|---|---|---|
| **+ 300 ppm; Control** | ∆SR | 0.09095 |

| | | | |
|---|---|---|---|
| **+ 300 ppm; Control** | ΔGini | 0.5015 | |
| **+ 150 ppm; Control** | ΔSR | 0.33095 | |
| **+ 150 ppm; Control** | ΔGini | 0.38675 | |
| **+ 150 ppm; + 300 ppm** | ΔSR | 0.21355 | |
| **+ 150 ppm; + 300 ppm** | ΔGini | 0.41055 | |
| **+ 1.5 °C; Control** | ΔSR | 0.2571 | |
| **+ 1.5 °C; Control** | ΔGini | 0.38985 | |
| **+ 3 °C; Control** | ΔSR | 0.4333 | |
| **+ 3 °C; Control** | ΔGini | 0.4144 | |
| **+ 3 °C; + 1.5 °C** | ΔSR | 0.35675 | |
| **+ 3 °C; + 1.5 °C** | ΔGini | 0.48675 | |
| **+ 150 ppm/+ 1.5 °C; Control** | ΔSR | 0.02295 | * |
| **+ 150 ppm/+ 1.5 °C; Control** | ΔGini | 0.4427083 | |
| **+ 300 ppm/+ 3 °C; Control** | ΔSR | 0.434 | |
| **+ 300 ppm/+ 3 °C; Control** | ΔGini | 0.3741667 | |
| **+ 300 ppm/+ 3 °C; + 150 ppm/+ 1.5 °C** | ΔSR | 0.0494 | * |
| **+ 300 ppm/+ 3 °C; + 150 ppm/+ 1.5 °C** | ΔGini | 0.3594583 | |
| **+ 300 ppm/+ 3 °C/drought; Control** | ΔSR | 0.439 | |
| **+ 300 ppm/+ 3 °C/drought; Control** | ΔGini | 0.4573 | |
| **Control/drought; Control** | ΔSR | 0.03805 | * |
| **Control/drought; Control** | ΔGini | 0.18865 | |
| **+ 300 ppm/+ 3 °C/drought; Control/drought** | ΔSR | 0.0341 | * |
| **+ 300 ppm/+ 3 °C/drought; Control/drought** | ΔGini | 0.2295 | |

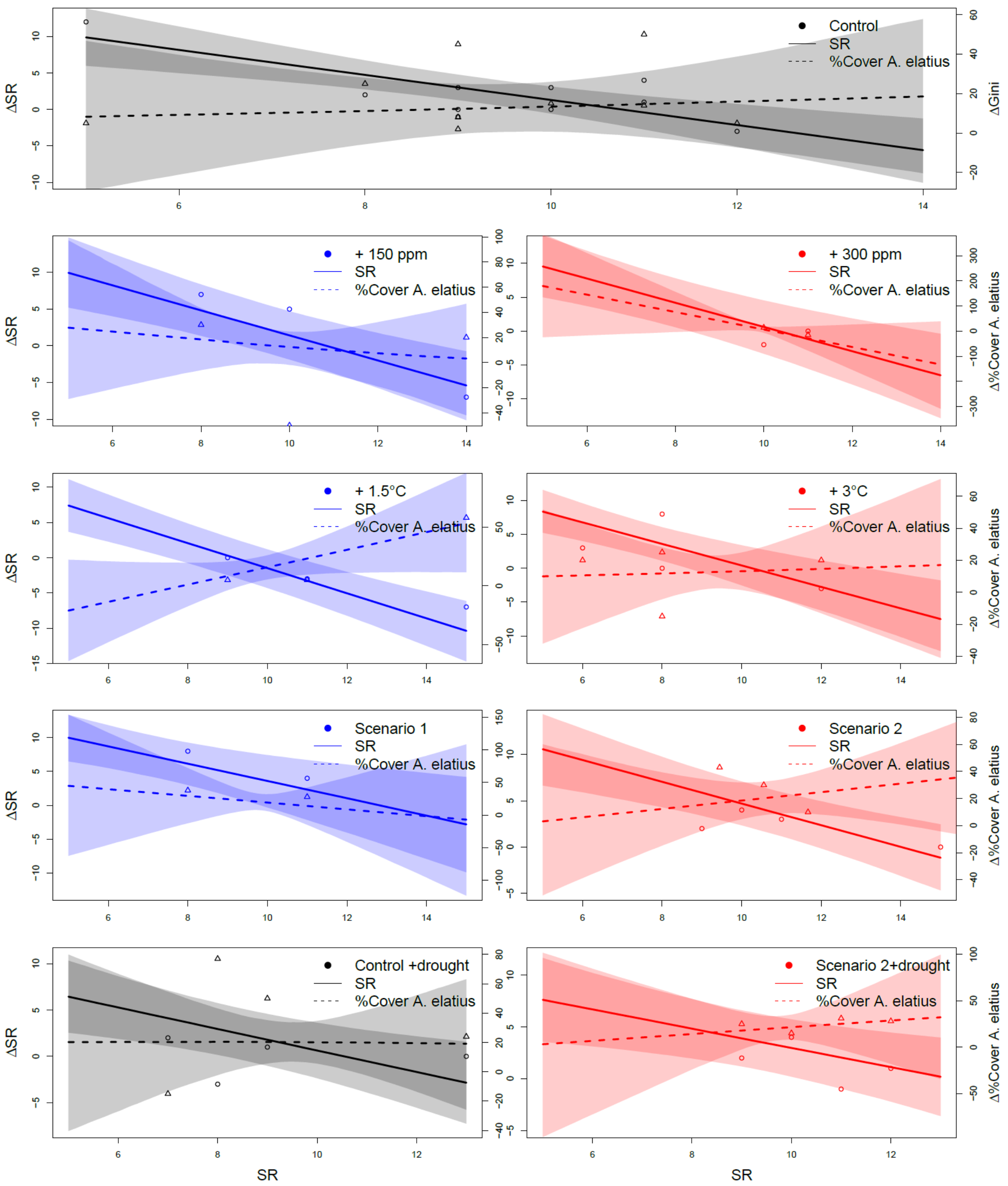

**Figure S5: brms models of the difference in species richness and Δ%cover *A. elatius* , before (2017) and after (2019) reseeding.** Regressions and confidence intervals are shown. Color indicates treatment levels in **a.** $CO^2$ treatments (+150 ppm and +300 ppm), **b.** temperature treatments (+1.5 °C and +3°C), **c.** scenario 1 (+150 ppm, + 1.5°C) and scenario 2 (+300 ppm and +3°C), and **d.** drought treatments (in control and scenario 2). P-values of difference between treatments can be found in **Tab. S5 and S6.**

**Table S5: p-values for differences in slopes of Δspecies richness and Δ%cover *A. elatius* (before and after reseeding) vs. species richness before reseeding, derived from brms posterior distributions**

| Comparison | type | p-value |
|---|---|---|
| **+ 300 ppm; Control** | $\Delta$%cover | 0.3812 |
| **+ 150 ppm; Control** | $\Delta$%cover | 0.3896 |
| **+ 150 ppm; + 300 ppm** | $\Delta$%cover | 0.3844 |
| **+ 1.5 °C; Control** | $\Delta$%cover | 0.4652 |
| **+ 3 °C; Control** | $\Delta$%cover | 0.47775 |
| **+ 3 °C; + 1.5 °C** | $\Delta$%cover | 0.49145 |
| **+ 150 ppm/+ 1.5 °C; Control** | $\Delta$%cover | 0.4862083 |
| **+ 300 ppm/+ 3 °C; Control** | $\Delta$%cover | 0.4975417 |
| **+ 300 ppm/+ 3 °C; + 150 ppm/+ 1.5 °C** | $\Delta$%cover | 0.484125 |
| **+ 300 ppm/+ 3 °C/drought; Control** | $\Delta$%cover | 0.4688 |
| **Control/drought; Control** | $\Delta$%cover | 0.4974 |
| **+ 300 ppm/+ 3 °C/drought; Control/drought** | $\Delta$%cover | 0.4696 |

**Table S6: Drought treatments** in the ClimGrass experiment

| | |
|---|---|
| 1. Drought experiment (2nd growth): | 23.05.2017 - 27.07.2017 |
| 2. Drought experiment (1st growth): | 18.04.2019 - 17.06.2019 |
| 3. Drought experiment (2nd growth): | 17.06.2020 - 29.07.2020 |
| 4. Drought experiment (2nd growth): | 27.05.2021 – 03.08.2021 |
| 5. Drought experiment (2nd growth): | 04.06.2022 – 29.07.2022 |
| 6. Drought experiment (2nd growth): | 22.05.2023 – 25.07.2023 |

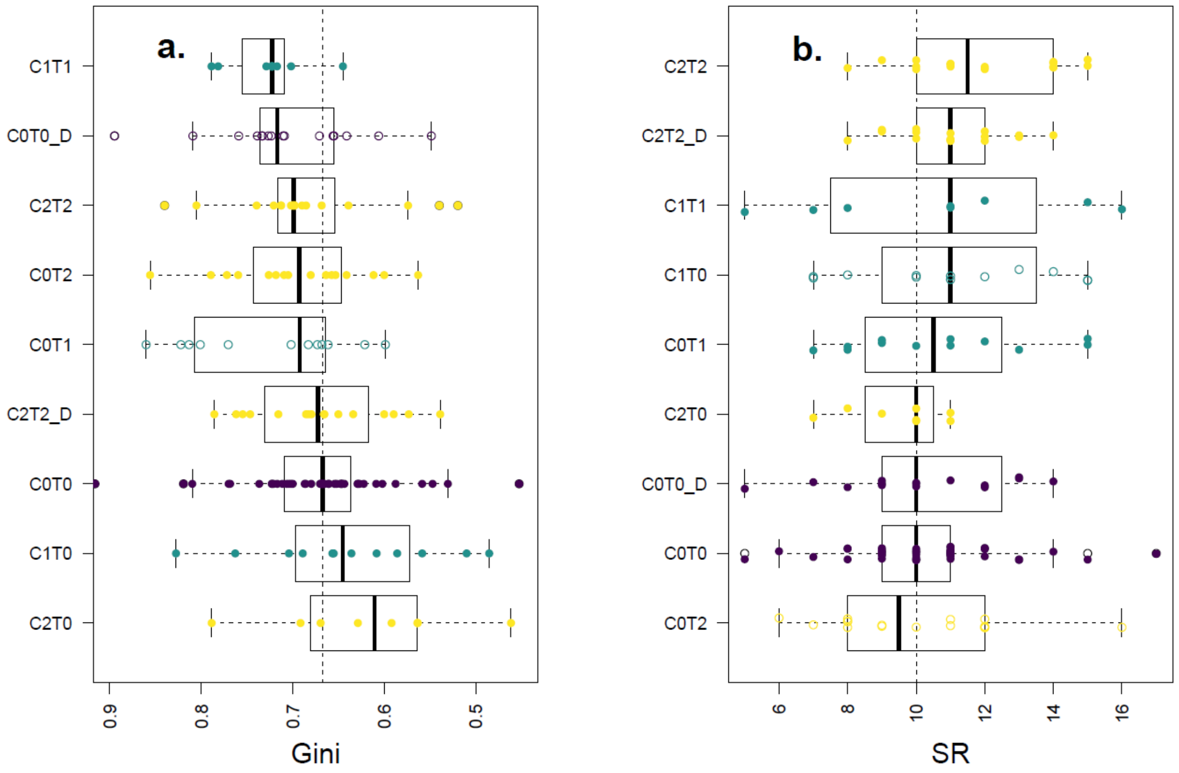

Fig. S6 **Treatment differences** in **a.** mean evenness, **b.** mean species richness. Data is included from all plots prior to reseeding in 2018, so only data before 2018 is included in the boxplots. Dashed line indicates value of the control (C0T0). Black bars indicate median, and boxes indicate 25% and 75% quantiles.  Treatment codes indicates temperature (T1 = +1.5 °C, T2 = +3°C) and $CO_2$ (C1 = +150 ppm, T2 = +300 ppm) levels and drought treatment (D = drought).

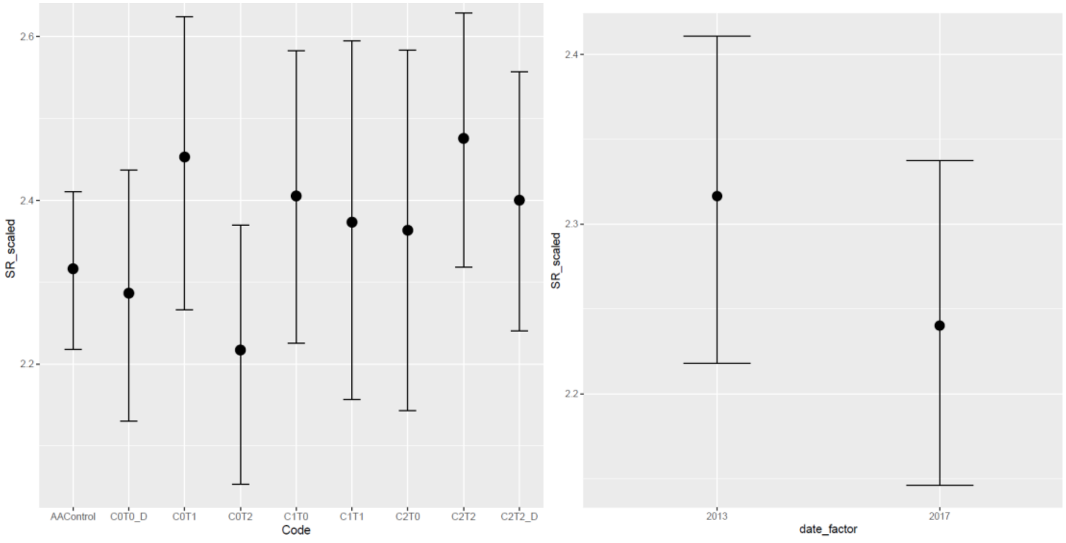

**Fig. S7** Difference in Species richness between treatments and between years. Included data is only from 2013 and 2017 from plots that were later in 2018 reseeded. Species richness (SR) on logarithmic scale. Plot is included as random effect; Year is included as categorial fixed effect (Formular: SR ~ Code + date_factor + (1|plot)).

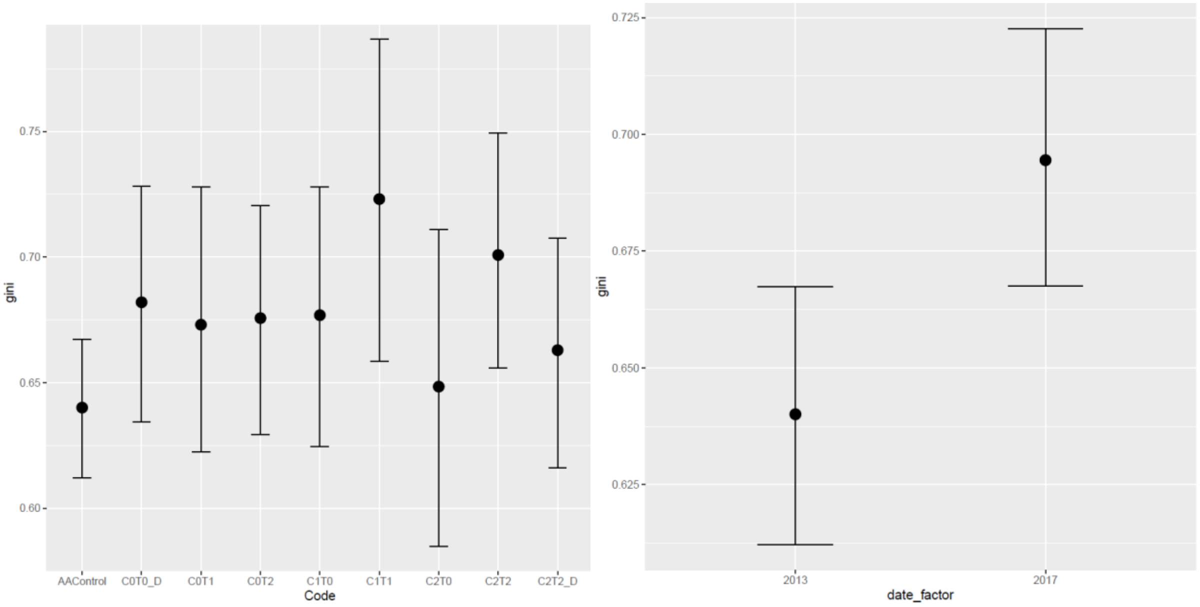

**Fig. S8** Difference in Evenness (Gini) between treatments and between years. Included data is only from 2013 and 2017 from plots that were later in 2018 reseeded. Plot is included as random effect; Year is included as categorial fixed effect (Formular: gini ~ Code + date_factor + (1|plot)).

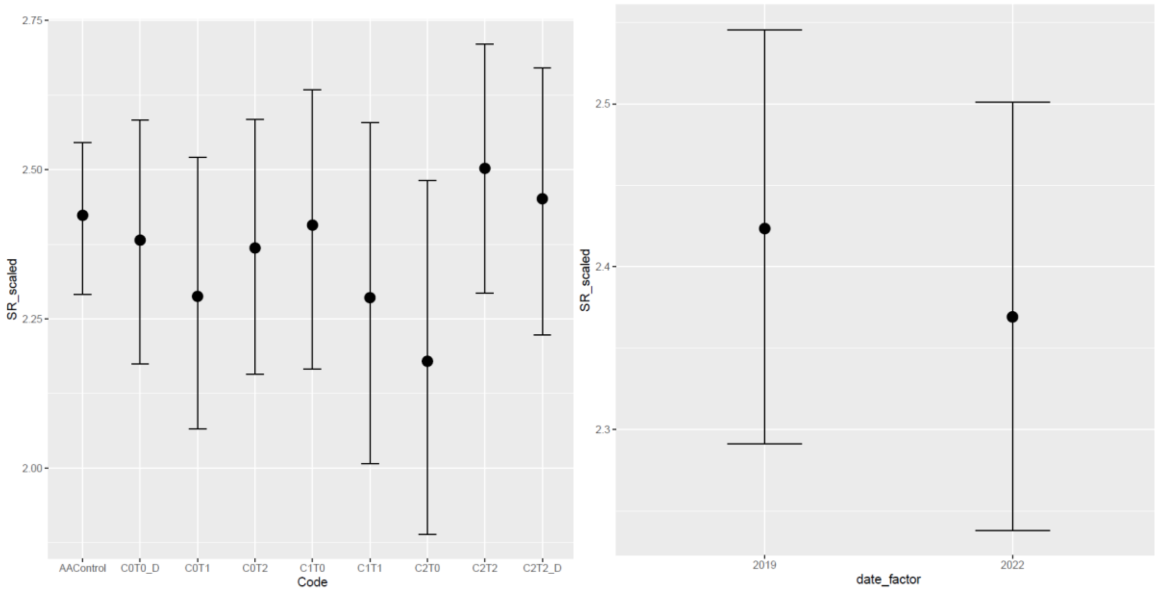

**Fig. S9** Difference in Species richness between treatments and between years. Included data is only from 2019 and 2022 from plots that were reseeded in 2018. Species richness (SR) on logarithmic scale. Plot is included as random effect; Year is included as categorial fixed effect (Formular: SR ~ Code + date_factor + (1|plot)).

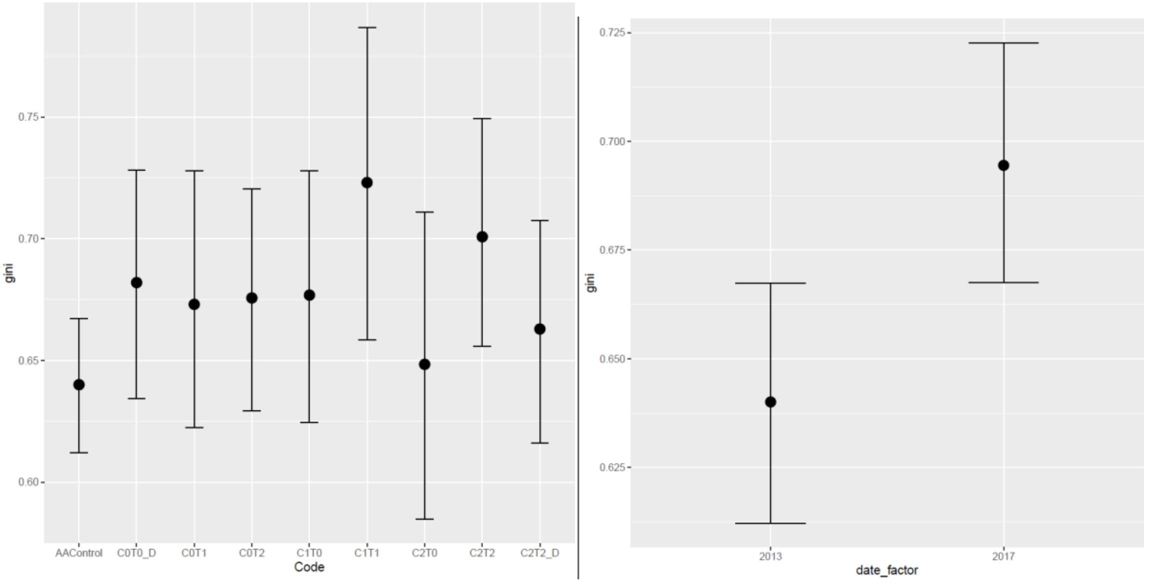

**Fig. S10** Difference in Evenness (Gini) between treatments and between years. Included data is only from 2013 and 2017 from plots that were later in 2018 reseeded. Plot is included as random effect; Year is included as categorial fixed effect (Formular: gini ~ Code + date_factor + (1|plot)).

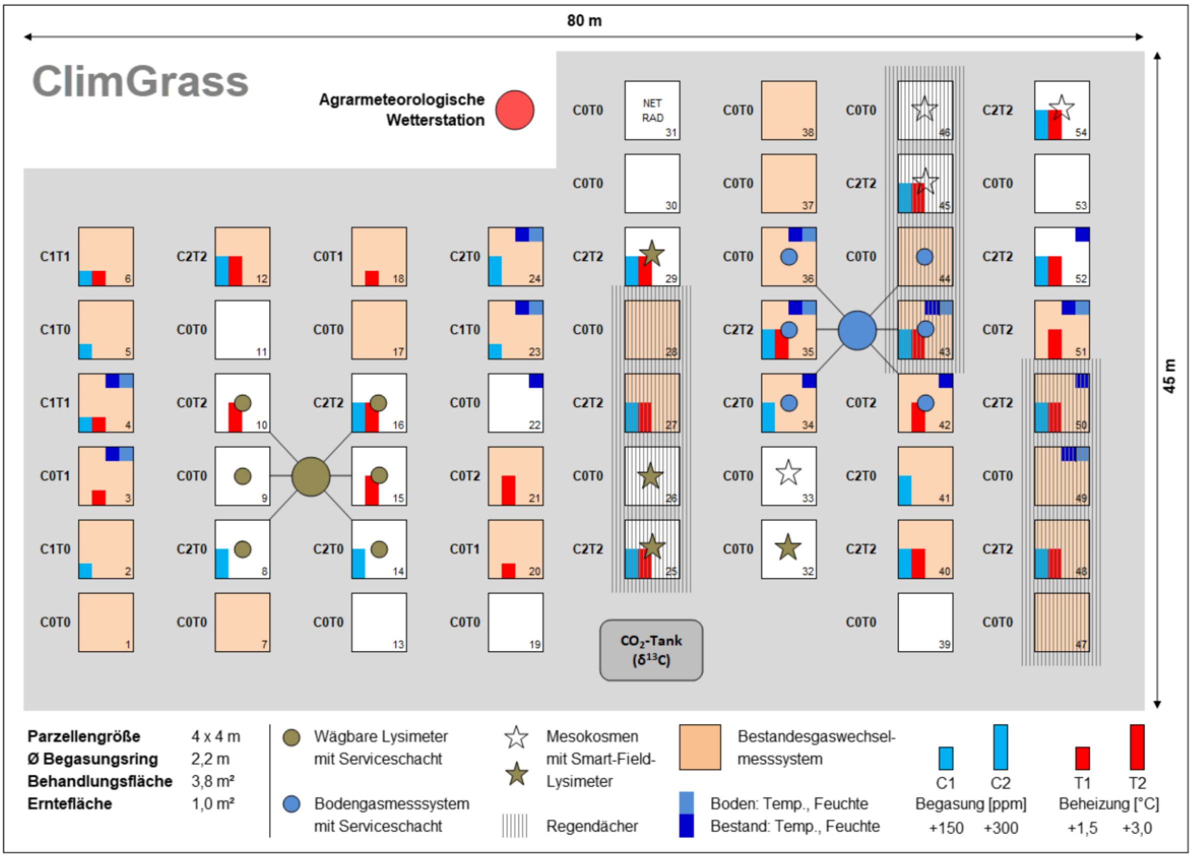

**Fig. S11**: Detailed set up of the ClimGrass experiment
